# CLN3 deficiency leads to neurological and metabolic perturbations during early development

**DOI:** 10.1101/2023.03.17.533107

**Authors:** Ursula Heins-Marroquin, Randolph R. Singh, Simon Perathoner, Floriane Gavotto, Carla Merino Ruiz, Myrto Patraskaki, Gemma Gomez-Giro, Felix Kleine Borgmann, Melanie Meyer, Anaïs Carpentier, Marc O. Warmoes, Christian Jäger, Michel Mittelbronn, Jens C. Schwamborn, Maria Lorena Cordero-Maldonado, Alexander D. Crawford, Emma L. Schymanski, Carole Linster

## Abstract

Juvenile Neuronal Ceroid Lipofuscinosis (or Batten disease) is an autosomal recessive, rare neurodegenerative disorder that affects mainly children above the age of 5 years and is most commonly caused by mutations in the highly conserved *CLN3* gene. Here, we generated *cln3* morphants and stable mutant lines in zebrafish. Although neither morphant nor mutant *cln3* larvae showed any obvious developmental or morphological defects, behavioral phenotyping of the mutant larvae revealed higher basal activity, hyposensitivity to abrupt light changes and hypersensitivity to pro-convulsive drugs. Importantly, in-depth metabolomics and lipidomics analyses revealed significant accumulation of several glycerophosphodiesters (GPDs) and a global decrease of bis(monoacylglycero)phosphate (BMP) species, two classes of molecules previously proposed as potential biomarkers for *CLN3* disease based on independent studies in other organisms. We could also demonstrate GPD accumulation in human-induced pluripotent stem cell-derived cerebral organoids carrying a pathogenic variant for *CLN3*. Our models revealed that GPDs accumulate at very early stages of life in the absence of functional CLN3 and highlight glycerophosphoinositol and BMP as promising biomarker candidates for pre-symptomatic *CLN3* disease.

## Introduction

Neuronal ceroid lipofuscinoses (NCLs) are a group of rare neurodegenerative lysosomal storage disorders mainly characterized by the accumulation of hydrophobic autofluorescent material in the lysosomes (Palmer et al. 1992; Dolisca et al. 2013; Carcel-Trullols, Kovacs, and Pearce 2015). To date, more than 10 genes (designated *CLN1-14*) have been linked to different forms of NCL. In particular, mutations in *CLN3* have been associated with a subclass of NCLs termed juvenile NCL (JNCL) (Dolisca et al. 2013; Jalanko and Braulke 2009) due to its onset between 5 to 8 years of life. JNCL is characterized by progressive vision loss, intellectual delay, behavioral changes, epileptic seizures in some cases, and premature death (Schulz et al. 2013). Given the high conservation at protein sequence level, models have been developed in several organisms to study the molecular and physiological functions of CLN3 and the mechanism underlying the associated disease (Cotman et al. 2002; Eliason et al. 2007; Gachet et al. 2005; Pearce et al. 1999).

The *CLN3* gene encodes a transmembrane protein of 438 amino acids localized in the endolysosomal compartment (Croopnick, Choi, and Mueller 1998; Ezaki et al. 2003) and, at the functional level, has been mainly associated with pH homeostasis, protein trafficking, calcium signaling, phospholipid distribution, and lysosomal amino acid transport (Gachet et al. 2005; Pearce et al. 1999; Padilla-Lopez et al. 2012; Chang et al. 2007). Despite extensive research efforts, the actual function of CLN3 in all these processes remains enigmatic and there is no effective treatment for *CLN3* patients that can prevent or at least slow down the disease progression. The main challenges for development of new therapies are the low number of patients available for enrolment in controlled clinical trials and the scarcity of patient-derived samples. To progress in disease understanding and therapy development despite these challenges, four different *Cln3* mutant mouse strains have been developed so far. They recapitulate the classical cellular hallmarks of JNCL, including intracellular accumulation of autofluorescent material and lysosomal dysfunction (Katz et al. 1999; Mitchison et al. 1999; Cotman et al. 2002; Eliason et al. 2007). All mouse models exhibit late onset of disease (6-12 months of age) characterized by slow progressive neurodegeneration, motor dysfunction, loss of specific neuronal populations, microglial activation, and astrocytosis. However, there are notable differences between the strains in terms of the onset of phenotypic abnormalities and the affected neuronal populations, indicating that genetic background and the genetic manipulation strategy employed can modulate the phenotypic outcomes in different *CLN3*-deficient models (Cotman et al. 2002; Mitchison et al. 1999; Katz et al. 1999; Kovacs and Pearce 2015).

Because of the high cost and relatively late disease onset of rodent models, it is highly relevant to develop other models to speed up *CLN3*-related research and address the urgent needs of the patients. Human iPSC-derived cerebral organoids homozygous for the CLN3^Q352X^ mutation were recently shown to recapitulate some of the NCL phenotypes (*i.e.* fingerprint-like intracellular inclusions, accumulation of subunit c of the mitochondrial ATP synthase (SCMAS), and lysosomal dysfunction) and represent thus a promising model to investigate the impact of *CLN3* deficiency on brain development and neuronal function. In the last years, zebrafish emerged as a highly relevant and convenient whole organism model to study developmental and mechanistic aspects of various human diseases (Santoriello and Zon 2012). Its external development and optical clarity allow for monitoring early development of organs such as the eye and the brain, which are among the most heavily affected organs in JNCL. A previous study using morpholino technology to knock down *cln3* in zebrafish resulted in morphant larvae showing embryonic dysmorphology, lysosomal dysfunction, epileptic seizures, and premature death (Wager et al. 2016). In the present study, we tested different non-overlapping morpholinos against the *cln3* gene and we could not detect any obvious phenotypic changes. Moreover, we created two stable *cln3* mutant zebrafish lines using the CRISPR/Cas9 technology and, in agreement with *CLN3* knockout models in other species, we confirmed that this gene is not essential for early organismal development. In fact, homozygous mutants reached adulthood without showing any gross phenotype and were indistinguishable in size and overall morphology from wildtype siblings. However, a closer analysis of the locomotor behavior revealed that *cln3* mutant larvae show higher basal activity and higher sensitivity to the effects of pro-convulsive drugs. Contrary to the relatively subtle behavioral changes, LC-MS-based metabolomics and lipidomics analyses revealed very interesting and striking differences between wildtype and *cln3* mutant larvae even at the very early stage of 5 days post-fertilization (dpf). Glycerophosphoinositol (GPI) emerged as the most significantly elevated metabolite in the *cln3* null mutant zebrafish larvae and bis(monoacylglycero)phosphate (BMP) 22:6 as one of the most significantly decreased lipid species. Importantly, we could demonstrate accumulation of certain glycerophosphodiesters (GPDs), including GPI, also in *CLN3* deficient cerebral organoids derived from human induced pluripotent stem cells (hiPSCs), further supporting that GPD accumulation might start very early during brain development in the absence of functional CLN3. In summary, *cln3-*deficient zebrafish do not develop detrimental neurological defects but display behavioral phenotypes as well as robust and specific metabolic perturbations, also observed in mammalian *CLN3*-deficient systems, at very early developmental stages. Our stable *cln3* mutant zebrafish lines represent therefore a novel tool to make rapid progress in elucidating the physiological function of CLN3 and potentially screen for molecules that can rescue CLN3 function and benefit Batten disease patients.

## Results

### CLN3 conservation and expression in zebrafish

We found that the zebrafish genome encodes one *CLN3* ortholog candidate by blasting the human CLN3 protein (NP_001035897.1) against the zebrafish (taxid:7955) proteome using the BLASTP 2.8.0 tool from NCBI. Only Battenin or Cln3 (NP_001007307.1), a protein of 446 amino acids, showed a significant alignment to human CLN3, with a sequence similarity and identity of 65% and 51%, respectively. Residues associated with Batten disease in human CLN3 are conserved in the zebrafish sequence, providing a first argument in favor of zebrafish being a relevant organism for modeling this disease (**Supplementary Fig. 1A**).

By analyzing *cln3* gene expression during embryonic development (6 hours post-fertilization (hpf) to 7 days post-fertilization (dpf)), we observed that *cln3* transcript levels increase during early developmental stages and stabilize after 4 dpf (**Supplementary Fig. 1B**). In addition, *cln3* transcript levels measured in various organs (brain, eye, heart, fin, skin, intestine, and gonads) of adult zebrafish females and males indicate ubiquitous expression with highest levels in the brain, eyes and gonads (**Supplementary Fig. 1C-D**). These observations only partially overlap with human and mouse expression data, in which *CLN3* is expressed at similar levels in all the analyzed organs (Yue et al. 2014; Fagerberg et al. 2014). Our results suggest that, in zebrafish, *cln3* might play an important role more specifically in the brain, eyes, and gonads during adulthood.

### Transient knockdown of *cln3* in zebrafish

Since *cln3* is expressed at very early stages of zebrafish development, we first attempted to knock down gene expression using morpholino oligonucleotides (MOs) as a rapid approach to analyze the impact of *cln3* deficiency as compared to generating stable knockout lines. We injected 8 ng of a translation-blocking MO (TB-MO) or splice-blocking MO (SB-MO) and monitored the overall embryonic morphology up to 7 dpf **(Fig. 1A-C)**. We used two different SB-MOs that were designed to target the boundaries between exon 2 and intron 2 (e2i2), and intron 2 and exon 3 (i2e3), respectively **(Fig. 1A)**. Based on PCR amplification of the targeted exon-intron boundaries and subsequent analysis by agarose gel electrophoresis, we could confirm that for both SB-MOs, the wildtype band (261 bp) disappeared to be replaced by novel bands whose lengths were consistent with *in silico* predictions based on exon skipping or intron inclusion **(Fig. 1D)**. Transcript sequence alterations were confirmed by sequencing to result in premature stop codons encoding early truncated proteins of 28-29 aa, assuming the transcripts are stable enough to be translated **(Fig. 1E)**. Due to the absence of reliable Cln3 antibodies, the efficiency of the MOs could not be confirmed at the protein level. All three *cln3* morphants were morphologically indistinguishable from their age-matched controls, as shown by representative individuals from the different groups at 3 dpf **(Fig. 1C)**. At higher morpholino concentrations, we observed toxicity in all embryos including those injected with the control morpholino. Altogether, the results strongly suggest that our knockdown approach is efficient and the absence of any obvious phenotype in the morphants indicates that *cln3* might not play an essential role during the early development in zebrafish.

**Figure 1.**
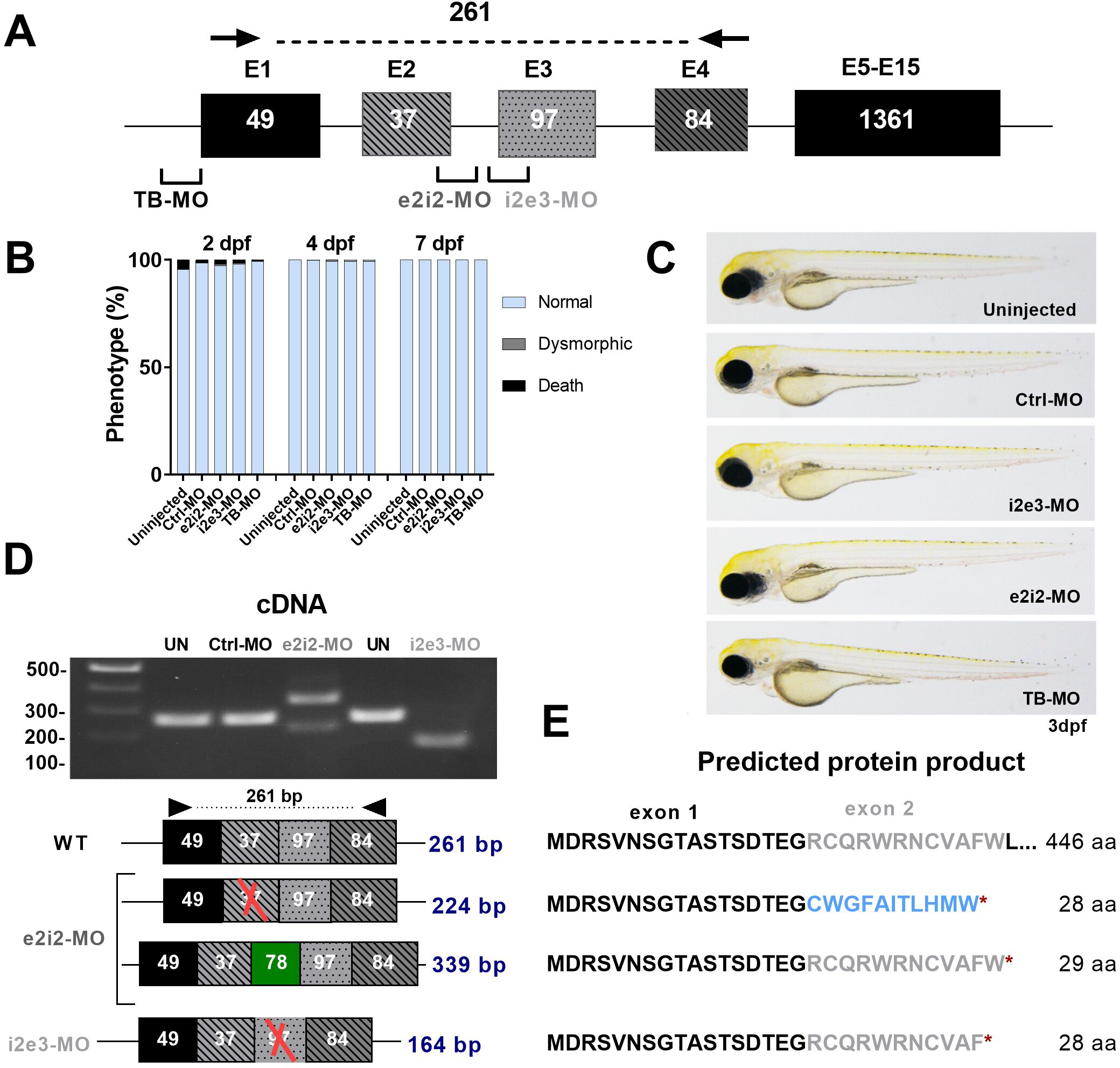
Transient knockdown of *cln3* in zebrafish larvae. **(A)** Schematic illustration of the *cln3* gene showing the target sites for the translation blocking (TB) and the two splice blocking (SB; i2e3-MO, e2i2-MO) morpholinos (MO) used in this study. Black arrows show the binding sites for the PCR primers used for SB-MO validation. **(B-C)** Microinjection of 8 ng SB- or TB-blocking MOs did not lead to any obvious morphological phenotype and larvae displayed a normal development. 150-200 larvae were analyzed per category at the indicated developmental time points. **(D)** Validation of MO efficiency by PCR amplification of target regions in cDNA of the *cln3* morphants in comparison to uninjected (UN) and control morpholino (Ctrl-MO) injected larvae. **(E)** Schematic representation of transcribed sequences and resulting translation products after treatment with the SB-blocking morpholinos. Microinjection of i2e3-MO resulted in exon 3 skipping, leading to an early stop codon. In contrast, e2i2-MO treatment led to two different splicing events, both also resulting in a premature stop codon. Excluded exons are marked with a red cross, included intron in green, and new amino acid sequence in light blue.

### Generation of stable *cln3* knockout lines using CRISPR/Cas9

Mouse models for *Cln3* deficiency showed a relatively late onset of symptoms (6-12 months), including several hallmarks of Batten disease such as neurodegeneration and lipofuscin accumulation (Cotman et al. 2002; Kitzmuller et al. 2008; Mitchison et al. 1999). Furthermore, in humans the disease only appears in childhood, suggesting that *CLN3* function may not be critical for morphogenesis or early development but rather may be involved in homeostasis, particularly in the brain, the organ most affected in the disease. Hence, we suspected that the potential phenotypes in zebrafish might also appear later in juvenile or adult individuals. Since morpholinos are typically efficient only until 5 dpf, studies in juvenile stages are not possible using morpholino-mediated approaches. Consequently, we generated two stable homozygous mutant zebrafish lines using the CRISPR/Cas9 technology by targeting the exon 4 – intron 4 boundary of the *cln3* gene (**Fig. 2A** and Supplementary Fig. 2A-C). The first line (*cln3^−/−^*; institutional code *cln3^lux1^,* hereafter referred to as MUT1), carries a long indel mutation that inserted 496 extra bp in the genomic DNA **(Fig. 2B)**. The insertion contains two copies of intron 4 and one copy of exon 5 **(Fig. 2C)**. Analysis of the cDNA revealed a main transcript encoding a truncated protein of 122 amino acids that contains 34 novel amino acids at the C-terminus **(Fig. 2D)** and only includes the first transmembrane domain **(Fig. 2E)**. This protein product is even shorter than the one described for the most common mutation (1.02 kb deletion) in JNCL patients (**Fig. 2E** and **Supplementary Fig. S3**). Therefore, we presume that the resulting protein has a complete loss-of-function and we considered the *cln3^lux1^* mutant as a knockout line. No change in *cln3* mRNA levels could be detected in MUT1 compared to wildtype larvae **(Supplementary Fig. 2F)**, indicating that the mutant transcript does not undergo nonsense-mediated mRNA decay.

**Figure 2.**
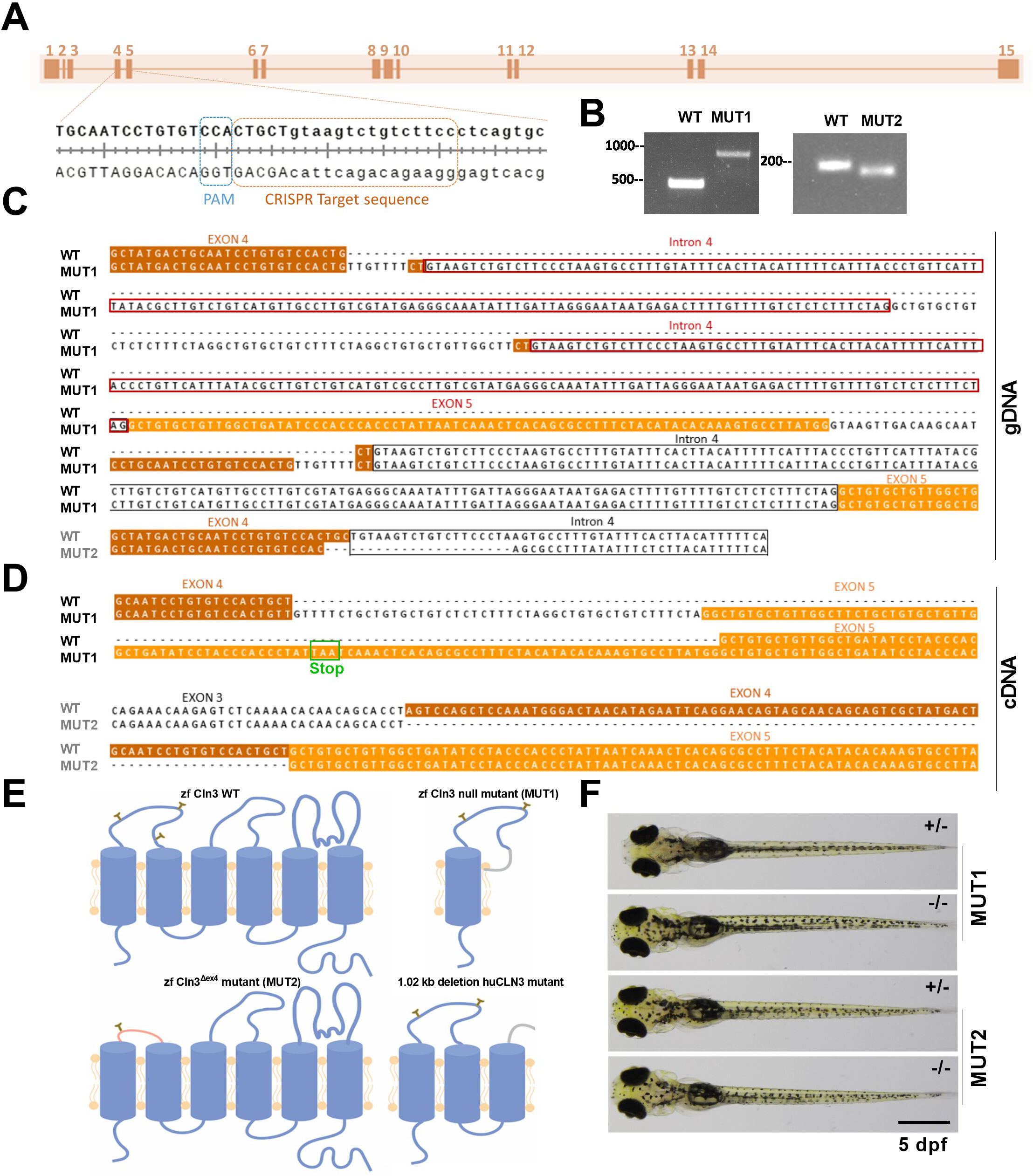
Generation of two stable *cln3* mutant lines in zebrafish using CRISPR/Cas9. **(A)** Schematic map of the *cln3* gene with zoom on the gRNA target site. Upper case letters represent the end of exon 4 and lower case letters the beginning of intron 4. **(B)** Gel electrophoresis of PCR amplicons from the mutated region in gDNA of F2 founders. **(C)** gDNA sequences of the two stable *cln3* mutant lines generated by CRISPR-Cas9 technology. **(D)** Based on cDNA sequence analysis, MUT1 carries an indel mutation resulting in a truncated translation product of 122 amino acids and MUT2 displays an in-frame deletion of exon 4. WT exon 4 sequence is highlighted in brown, exon 5 in orange, and intron 4 with a black box. Additional copies of intron 4 are highlighted with red boxes and the premature stop codon with a green box. **(E)** Schematic representation of the expected protein products encoded by the WT *cln3*, *cln3* null (MUT1), *cln3^Δex4^* (MUT2) alleles and the most common human Batten disease *CLN3* allele (1.02kb deletion; *huCLN3*) based on the predicted membrane topology described in (Mirza et al. 2019). **(F)** MUT1 and MUT2 mutants and heterozygous controls at 5dpf. Homozygous mutants are morphologically indistinguishable from their heterozygous counterparts. WT, wildtype; zf, zebrafish; hu, human.

The second mutant line (*cln3^Δex4/Δex4^*; institutional code *cln3^lux2^*, hereafter referred to as MUT2) has a deletion of 22 bp at the genomic DNA level **(Fig. 2B, C)** that disrupts the splicing process, resulting in an in-frame skipping of exon 4 at the cDNA level (first luminal loop of the transmembrane protein) and a predicted expression product that may retain residual function **(Fig. 2C-E)**. However, in both the human and zebrafish genes, exon 4 encodes two putative asparaginyl glycosylation sites (**Supplementary Fig. 3**) and contains a conserved serine residue mutated in some patients affected by Batten disease (Mirza et al. 2019). Therefore, we decided to proceed with both mutant lines to study phenotypic impacts that may be relevant for human *CLN3* disease.

To determine whether homozygous *cln3* mutants are viable, we raised the progeny from F1 incrosses to adulthood and genotyped the fish after 2 months. For both the null and exon 4 deletion mutations, 30-46% of the F2 progeny were homozygous mutants (Supplementary Fig. 2D), suggesting that *cln3* deficiency does not induce lethality at embryonic or juvenile developmental stages in zebrafish. Furthermore, homozygous mutants did not exhibit obvious differences in size, gross morphology, or fertility in comparison to their wildtype and heterozygous siblings. Finally, survival was not affected in homozygous MUT1 fish compared to their heterozygous siblings (**Supplementary Fig. 2E**).

### Behavioral assays in *cln3* mutant zebrafish larvae indicate early neurological defects

In zebrafish, the innate behavioral response depends on the correct development and functioning of the nervous system (Basnet et al. 2019). To uncover potential neurological impairments in the *cln3* mutant larvae, we therefore performed three behavioral assays: global activity, dark-light response, and pro-convulsive drug response **(Fig. 3A)**. In the first approach, we studied the global activity of larvae in dark or light condition, after a 15-min pre-adaptation period in the corresponding condition. In both conditions, MUT1 larvae showed a higher locomotor activity in comparison to heterozygous control larvae; for MUT2 larvae, a (non-statistically significant) higher basal activity was only observed in the light **(Fig. 3B-C)**. In the second approach, the lighting conditions were alternated in 15-min intervals (dark-light-dark), after a 15 min pre-adaptation period in the dark. In the first dark period, control larvae showed a constant velocity. During the light period, activity gradually increased, and in the subsequent dark period, larvae remained transiently hyperactive before recovering a more passive swimming mode **(Fig. 3D)**. Mutant larvae displayed a similar dark-light behavioral response, except during the dark-to-light and light-to-dark switches (approx. min 15-20 and 30-32). During these transition periods, the control larvae showed abrupt changes in activity, with notably a rather dramatic freeze startle response upon turning on the lights, which was virtually absent in both the MUT1 and MUT2 larvae. These results suggest that *cln3* deficient larvae, although capable to react to light changes, may have an impaired visual acuity.

**Figure 3.**
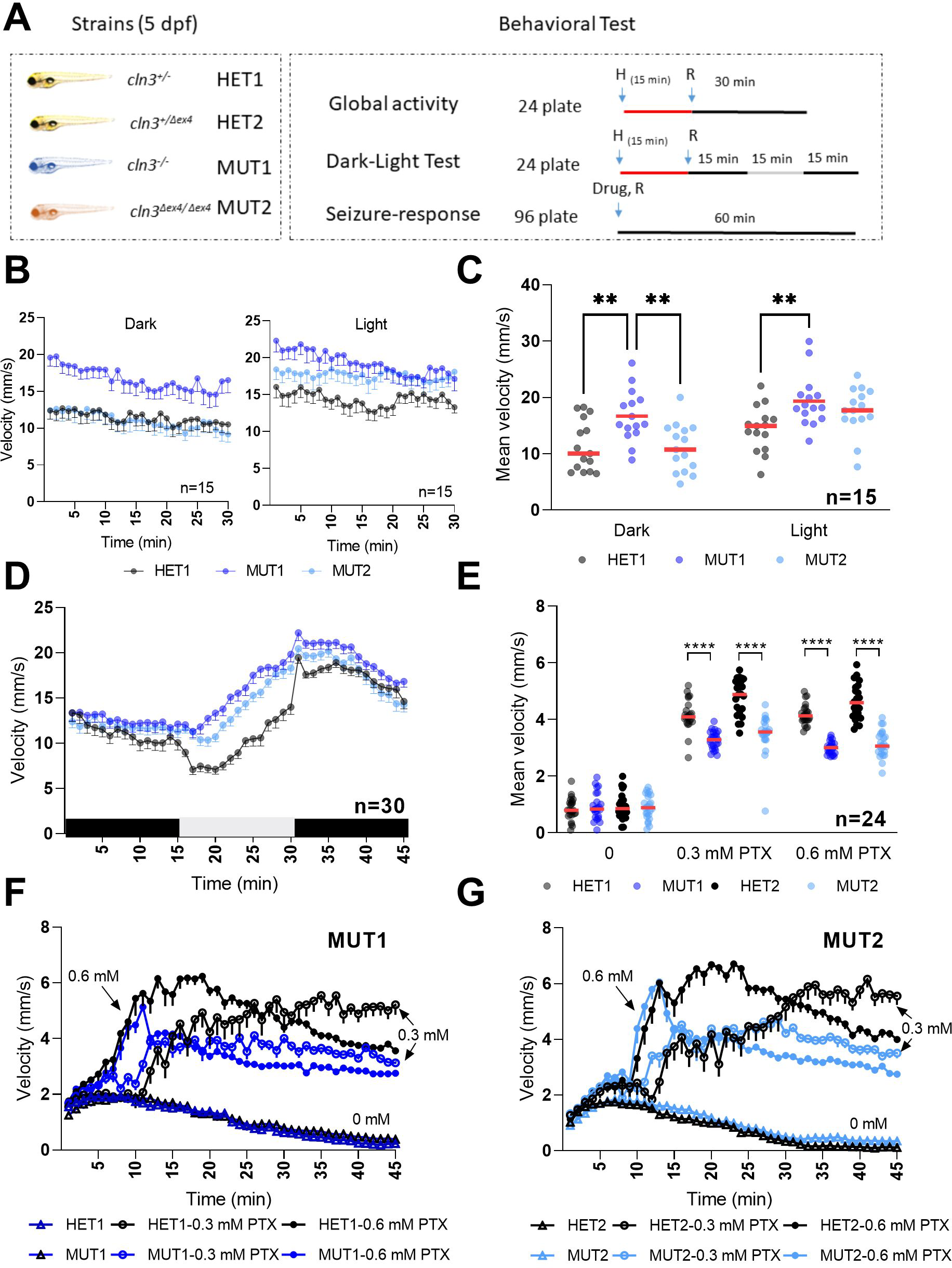
Locomotor behavior of *cln3* mutant larvae under different light conditions. **(A)** Experimental design of behavioral response studies. For the global activity and dark-light response assays, 8 larvae/line were placed in a 24-well plate (1 larva/well) and pre-adapted for 15 min to the indicated condition (light/dark) before starting the recording. The movement was tracked for 45 min in 15-min light/dark intervals or for 30 min under one continuous lightening condition. For the seizure-response assay, 12 larvae/line were placed in a 96-well plate and recording was started shortly after addition of PTX for 1 h. Vehicle control was 0.6% DMSO. Each experiment was performed at least in duplicate with 5 dpf larvae from different parental batches. **(B)** Mean velocity (1 min intervals) of the larvae recorded under constant lighting conditions. **(C)** Total mean velocity recorded in panel B. **(D)** Mean velocity (1 min intervals) of the larvae in alternating lighting conditions. **(E)** Total mean velocity in 1 hr exposure to 0, 0.3 and 0.6 mM PTX. **(F, G)** Behavioral profiles (1 min intervals) of the larvae recorded in panel E. In all the behavioral profiles shown, data points are means ± SEMs. In the scatter dot plots, each dot represents the mean velocity of one larva and the red line represents the median. Statistically significant differences between lines were determined using the ordinary two-way ANOVA test followed by Sidak’s multiple comparisons test (**, p ≤0.0021; ****, p ≤0.0001).

For the drug response assay, we exposed *cln3* mutant larvae to pentylenetetrazole and picrotoxin (PTZ and PTX, respectively). Both compounds are chemoconvulsant drugs that induce epileptic seizures in murine and zebrafish models (Yang et al. 2017; Kruse and Kuch 1985). First, MUT1 and heterozygous control larvae at 5 dpf (n=12) were incubated with different concentrations of PTZ (5-20 mM) under dark conditions. After 10 min habituation in the dark, the locomotor activity was recorded for 1 hr in that same condition. All tested PTZ concentrations induced an increase of activity in both the heterozygous and homozygous mutant *cln3* lines (**Supplementary Fig. 4A, B**). At subacute PTZ concentrations (7.5 and 10 mM), MUT1 larvae showed a significantly higher activity (2.01 ± 0.54 and 2.51 ± 0.39 mm/s) compared to the heterozygous control larvae (1.48 ± 0.34 and 1.94 ± 0.37 mm/s; values represent mean velocities moved during the 60 min). However, at acute PTZ concentrations (15 and 20 mM), total mean velocities between control and MUT1 larvae were not significantly different (**Supplementary Fig. 4A**).

PTX was tested at 0.01, 0.03, 0.1, 0.3, and 1 mM using the same protocol as applied for the PTZ treatment (**Supplementary Fig. 4C**). A significant increase of locomotor activity was detected starting only at 0.3 mM PTX in the control heterozygous larvae, whereas 0.1 mM PTX already induced a significantly higher activity in MUT1 larvae. In contrast to the results obtained with PTZ and unexpectedly, at 0.3 and 1 mM PTX, MUT1 larvae showed a remarkably lower movement response (2.82 ± 0.53 and 2.13 ± 0.23 mm/s) compared to the heterozygous control larvae (4.32 ± 0.52 and 3.18 ± 0.62 mm/s) (**Supplementary Fig. 4C, D**).

The PTX phenotype was confirmed in further experiments that included also the MUT2 zebrafish line **(Fig. 3E-G)**. All larvae showed a similar behavioral profile in vehicle condition (DMSO) (**Fig. 3F, G**). In the presence of PTX, control and mutant larvae displayed a sharp increase in activity within 5-10 min. However, the activity of the *cln3* mutants plateaued at a significantly lower level after 10-20 min, compared to the heterozygous control larvae. In conclusion, these results showed that *cln3* deficiency does not lead to spontaneous epileptic seizures, at least in these early stages of zebrafish larval development. However, the behavioral phenotypes observed in the presence of different concentrations of proconvulsant drugs clearly indicated that *cln3* is important for healthy neuronal function and that some mild neurological impairments can already be observed in young (5 dpf) larvae.

### Metabolic perturbations in *cln3* mutant larvae detected by untargeted metabolomics

Since we observed increased locomotor activity in the MUT1 line under normal conditions, we performed HILIC-HRMS-based untargeted metabolomics analysis in polar extracts of six pools of 40 MUT1 and 40 WT larvae at 5 dpf, obtained from different parental batches **(Fig. 4A)**. In this discovery set of samples, we detected 3553 features (*i.e.* peaks) in positive mode and 2775 features in negative mode (6328 in total) within the range of 60-900 *m/z* (**Supplementary Table 1**). Using a Student’s t-test, we found that 29% of the features (*i.e.* 1857) had differential abundances (p<0.05) between both lines. A total of 798 features (603 annotated) were lower in the mutant compared to WT larvae while 1059 features (766 annotated) were higher (**Supplementary Table 1** and **Fig. 4B**). Principal Component Analysis (PCA) using all measured features revealed a clear and consistent separation between WT and MUT1 samples **(Supplementary Fig. 5A)** suggesting profound differences at the metabolomic level. Lower clustering in this analysis for the MUT1 samples as compared to the WT samples, suggested a certain degree of metabolic divergence in the mutant genotype.

**Figure 4.**
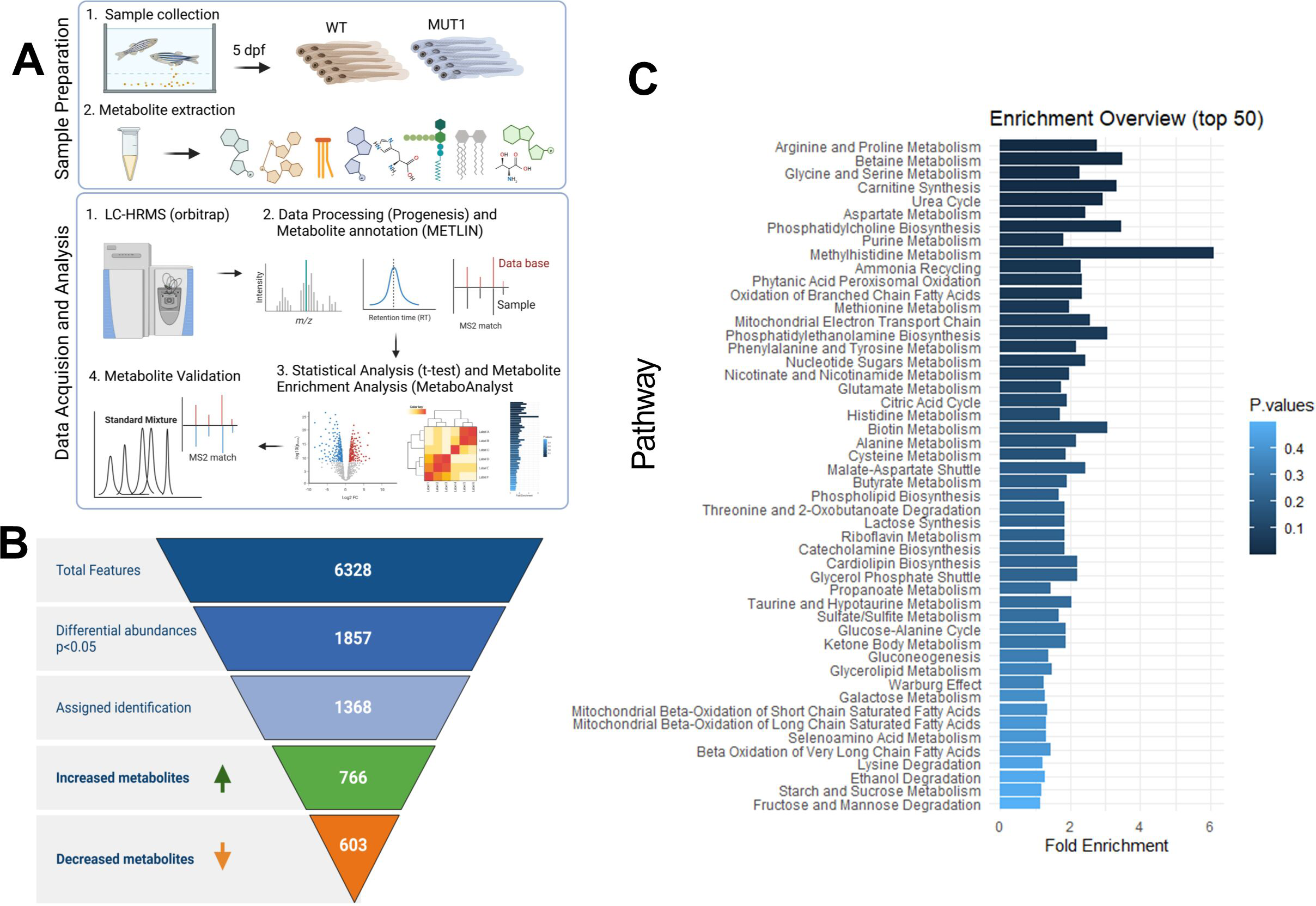
Untargeted metabolomics differentiates WT and MUT1 samples. **(A)** Experimental workflow of the untargeted metabolomics study from sample collection to data analysis for the discovery set of samples. **(B)** MS feature filtering pipeline including statistical analyses and compound identification. Out of 6328 detected features, 1857 differed significantly between the WT and MUT1 lines and, of those, 1369 could be annotated. **(C)** Metabolite enrichment analysis of differential metabolites (p<0.05) highlights the major dysregulated pathways between WT and MUT1 samples in the discovery batch.

Based on the metabolite enrichment analysis performed using MetaboAnalystR, carnitine synthesis, urea cycle and several amino acid pathways (*e.g*. “Arginine and Proline Metabolism”, “Glycine and Serine Metabolism” and “Aspartate Metabolism”) were ranked among the top perturbed metabolic pathways **(Fig. 4C)**. In addition, phosphatidylcholine biosynthesis, mitochondrial β-oxidation and TCA cycle were identified as being significantly altered in MUT1 larvae. All these pathways have been previously highlighted in studies using different NCL models and patient-derived cells, supporting the disease relevance of our zebrafish model (Dawson et al. 1996; Padilla-Lopez et al. 2012; Gomez-Giro et al. 2019; Chan, Ramirez-Montealegre, and Pearce 2009; Kim et al. 2012).

The most significantly changed metabolite in our comparative metabolomics analysis between WT and MUT1 larvae was glycerophosphoinositol (GPI), reaching ≥19-fold higher levels in the mutant extracts **(Fig. 5A, B)**. Adenosine diphosphate ribose was identified as the most decreased metabolite in MUT1 and glucosamine-6-phosphate and 1-alpha-D-galactosyl-sn-glycerol 3-phosphate were ranked as the (annotated) metabolites with the highest -fold change increase in MUT1. Interestingly, we could also observe a significant accumulation in other glycerophosphodiesters (GPDs) such as glycerophosphoglycerol (GPG; 1.4-fold) and glycerophosphocholine (GPC; 3-fold) **(Fig. 5B)**, whereas glycerophosphoethanolamine (GPE) was not significantly changed and glycerophosphoserine (GPS) was not detected in the extracts. While this work was ongoing, accumulation of GPDs has also been found in yeast and mammalian models of *CLN3* deficiency (Laqtom et al. 2022; Brudvig et al. 2022), suggesting that CLN3 plays a highly conserved role in maintaining normal GPD levels across species. Phospholipid precursors such as phosphocholine and phosphoethanolamine were also detected at significantly higher levels in the mutant compared to WT larvae **(Fig. 5B)**, reminiscent of a report of impaired phosphatidylethanolamine and phosphatidylcholine synthesis through the Kennedy pathway for *Saccharomyces cerevisiae* strains deficient in the CLN3 homologous protein battenin or BTN1 (Padilla-Lopez et al, 2012). Consistent with the metabolite enrichment analysis, we observed a significant decrease in the levels of certain amino acids in the MUT1 larvae, including aspartic acid, glutamine, arginine, ornithine, and proline **(Fig. 5C)**. Furthermore, some amino acid-derived neurotransmitters appeared to be dysregulated in MUT1 mutants **(Fig. 5D)**. The levels of N-acetylaspartylglutamic acid (NAAG), the third most abundant neurotransmitter in human brain and previously proposed as a biomarker of severity in *CLN2* disease (Sindelar et al. 2018), were decreased about 2-fold in MUT1 larvae compared to WT (p<0.0001). The inhibitory neurotransmitter γ-aminobutyric acid (GABA) unexpectedly showed slightly (10%) but significantly (p<0.01) higher levels in MUT1 larvae. Interestingly, norepinephrine sulphate reached detectable levels in MUT1, but not in WT extracts. We also observed alterations in metabolic pathways that are not present in the Small Molecule Pathway Database (SMPD) used for our metabolite enrichment analysis. Notably, acylcarnitine (AC) synthesis appeared to be perturbed, with significantly decreased levels measured for most ACs detected in MUT1 larvae, while the level of acetylcarnitine was slightly increased (**Fig. 5E, Supplementary Fig. 5A**).

**Figure 5.**
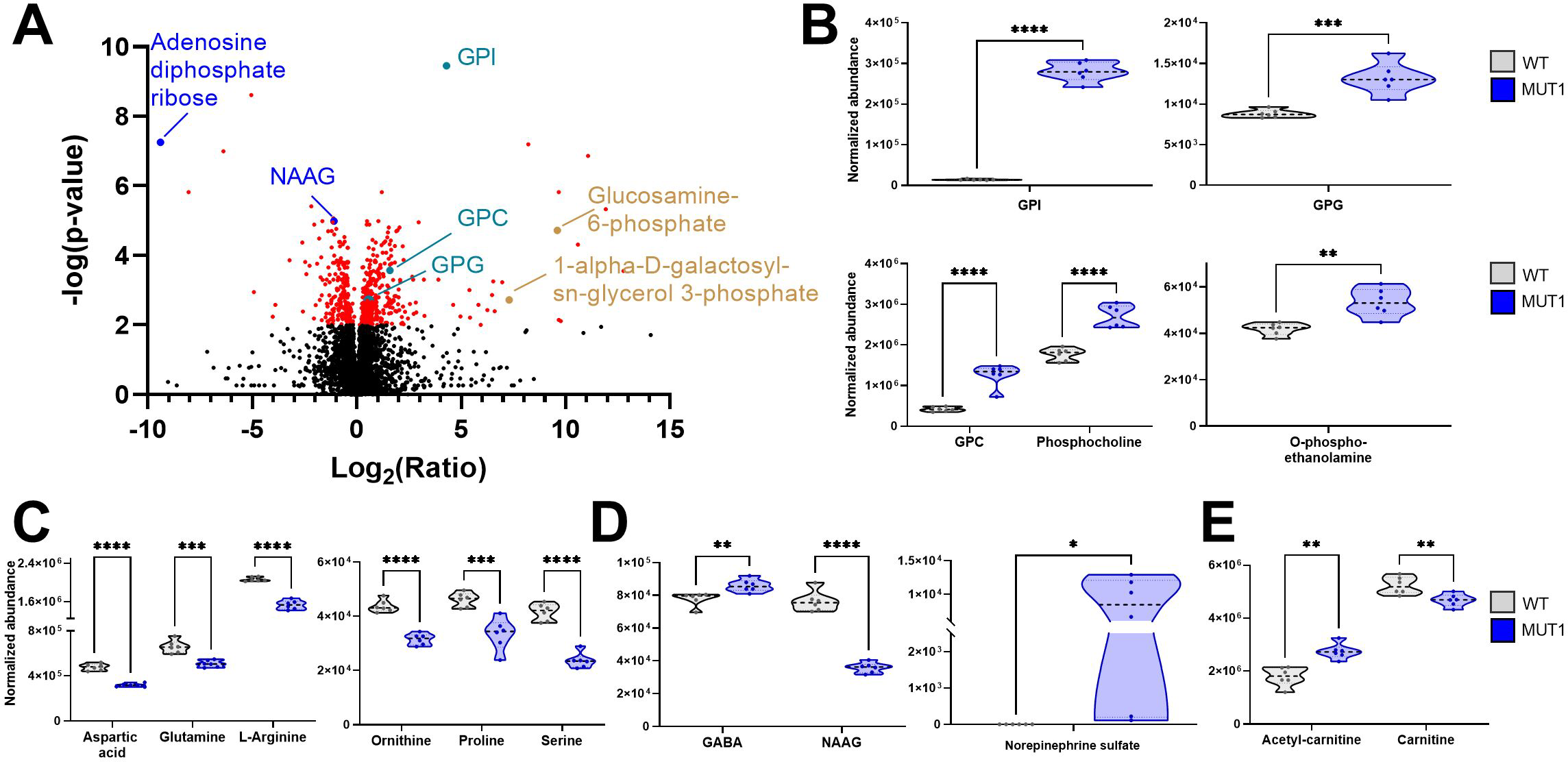
Most significantly altered metabolites between WT and MUT1 larvae based on untargeted metabolomics. **(A)** Volcano plot (corrected p-value vs. -fold change ratio) showing significantly (p < 0.001) altered metabolites in red. Each dot represents one metabolite. Adenosine diphosphate ribose (the most decreased metabolite in MUT1) and NAAG (previously reported as potential NCL biomarker (Sindelar et al. 2018)) are highlighted in blue and GPD species in turquoise. Metabolites with the highest -fold change among the ones with compound annotation are highlighted in brown. (**B-E**) Violin plots showing levels of selected differential metabolites as normalized abundance values. Statistically significant differences between the WT and MUT1 lines were determined using an unpaired multiple Welch’s t-test (*, p ≤ 0.05; **, p ≤ 0.01; ***, p ≤ 0.001; ****, p < 0.0001).

### Validation of metabolic disruptions and assessment of lysosomal proteolytic activity in MUT1 zebrafish larvae

An independent validation set of samples obtained from different parental zebrafish pairs was prepared (n=5). The experimental and data analysis workflows were identical to those described above for the discovery sample set. We detected 2795 features in positive and 3578 features in negative mode, with a total of 1391 features showing differential abundances (p<0.05 based on a Student’s *t*-test) in MUT1 zebrafish larvae compared to WT larvae (**Supplementary Table 2**, **Supplementary Fig. 6A**). In this data set, 760 features (of which 636 were annotated) were significantly higher and 631 features (of which 489 were annotated) were significantly lower in the mutant larvae (**Supplementary Fig. 6A**). As for our initial experiment, PCA analysis using all features confirmed the segregation between the WT and MUT1 lines (**Supplementary Fig. 6B**). Overall, metabolite enrichment analysis of the validation set corroborated the most significant pathway perturbations (amino acid metabolism, carnitine synthesis, urea cycle, phospholipid metabolism, and mitochondrial β-oxidation) found in the discovery set (**Supplementary Fig. 6C, Supplementary Fig. 7**).

Since we observed a reduction of proline levels in *cln3* mutants, we examinated our differentially abundant metabolites for oligopeptides containing prolyl residues. We observed increased levels of these oligopeptide species in the MUT1 extracts in both of our data sets (**Supplementary Fig. 8A**), which, in combination with the decreased levels of free amino acids, may suggest a decreased lysosomal proteolytic activity in MUT1 larvae. As one of the most common cellular phenotypes observed in Batten disease is decreased cathepsin D (CtsD) activity, we assessed the latter in whole larvae extracts. Although not statistically significant, we consistently measured lower CtsD activity in *cln3* mutant extracts, indicating lysosomal impairments that might be too mild to be detected at peptidase level by our assay at this developmental stage (**Supplementary Fig. 8B**). CtsD protein levels were similar between the control and mutant extracts (**Supplementary Fig. 8C**). Finally, transmission electron microscopy (TEM) in liver, intestine and muscle tissue of 5 dpf MUT1 larvae did not reveal characteristic differences at the cellular level such as fingerprint-like inclusions, suggesting that this phenotype is not evident at early developmental stages, but should be further investigated in adult tissues (**Supplementary Fig. 8D**).

The identity of 16 statistically significant differential metabolites (in both the discovery and validation sets) was confirmed by direct comparison with the retention time and MS2 spectrum of reference standards (**Supplementary Table 3**). We could not verify the identity of norepinephrine sulfate due to lack of the corresponding reference standard. As further validation of the untargeted metabolomics analysis, we performed targeted LC-MS-based measurements of GPDs, amino acids and ACs. New extracts were prepared from a new batch of larvae obtained from a different generation, allowing also to assess the conservation of the metabolic phenotypes across generations (**Fig. 6A, B**, **Supplementary Fig. 9A, B**). In full agreement with the untargeted results, we could confirm the accumulation of GPI, GPC, GPG and GPS in MUT1, as well as unchanged levels for GPE **(Fig. 6A, B)**. The GPI peak was barely detectable in WT samples (**Supplementary Fig. 9A**). Interestingly, using our targeted LC-MS/MS method, we did not find accumulation of GPDs in a zebrafish model knocked out for ATP13A2/CLN12 (previously described in (Heins-Marroquin et al. 2019)), suggesting that the abnormal GPD levels might be specific for CLN3 loss-of-function and not generalizable to all other NCL genes (**Supplementary Fig. 9B**). We also measured GPDs in hiPSC-derived cerebral organoids, a state-of-the-art human model for brain development previously described in (Gomez-Giro et al. 2019; Lancaster and Knoblich 2014). Strikingly, GPI and GPG levels were 3-to 10-fold higher in organoids carrying the pathogenic CLN3^Q352X^ variant compared to isogenic control organoids (**Fig. 6C**). We also found an increase in GPS, but not in GPE and GPC levels in the mutant organoids. Taken together, our results strongly indicate that GPDs accumulate very early during (brain) development under *CLN3* deficiency and that the role of CLN3 in maintaining normal GPD levels is conserved from fish to humans.

**Figure 6.**
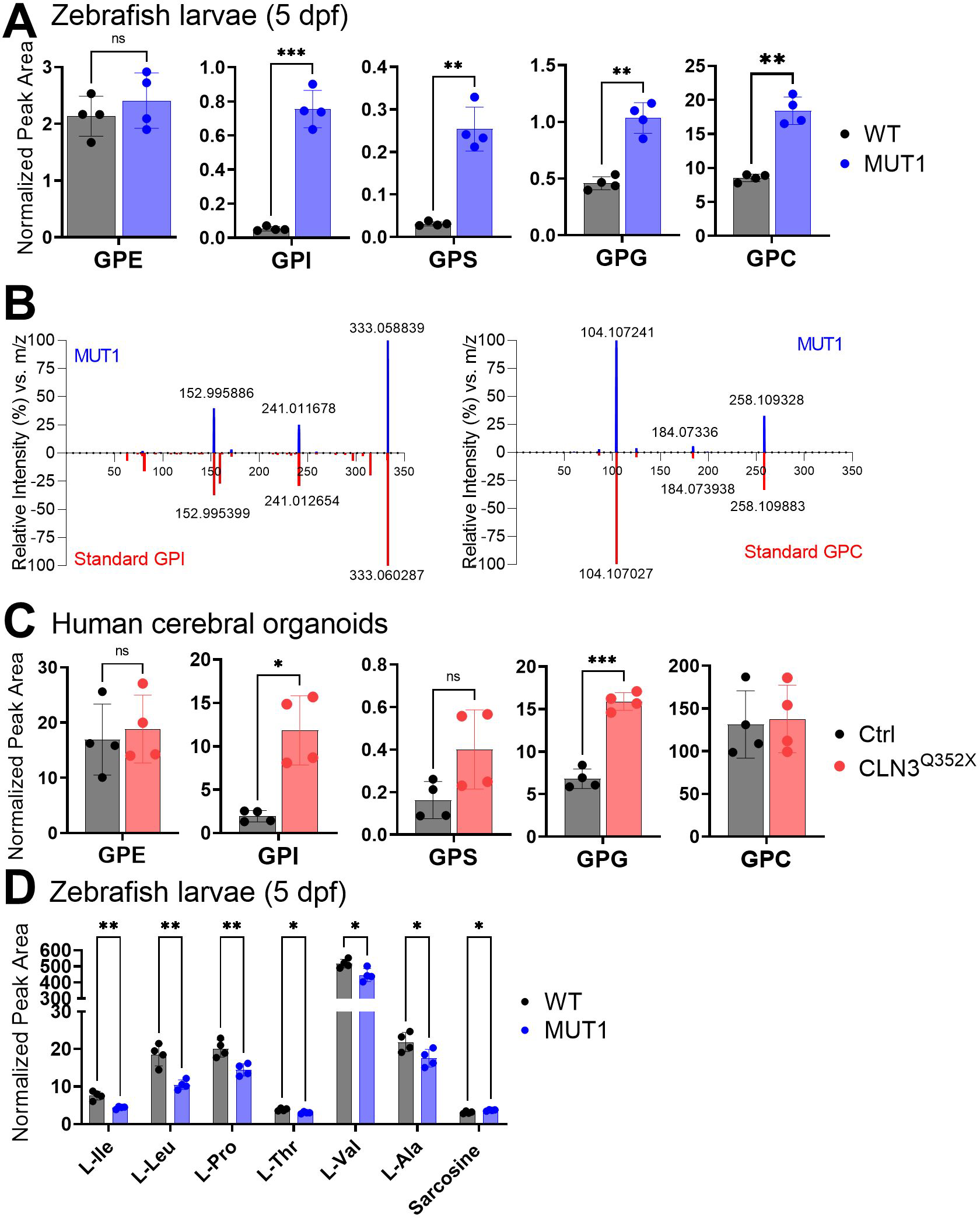
Validation of glycerophosphodiester and amino acid changes by targeted LC-MS analyses. **(A)** Metabolite levels of glycerophosphodiester species (GPE, GPI, GPS, GPG, GPC) extracted from 40 whole zebrafish larvae at 5 dpf. **(B)** Representative MS/MS spectrum plots of GPI and GPC from MUT1 extracts (in blue) and metabolite standards (in red), further confirming compound identities (matching library score 83%, isotopic pattern score 100%). **(C)** Metabolite levels of the same glycerophosphodiester species as shown in panel A, but extracted from human iPSC-derived cerebral organoids after 55 days of differentiation. **(D)** Metabolite levels of amino acids differing significantly between WT and MUT1 zebrafish larvae at 5 dpf based on targeted LC-MS measurements. Data shown are means ± SDs of four biological replicates. Statistically significant differences between WT and mutant samples were determined using an unpaired t-test (*, p ≤ 0.05; **, p ≤ 0.01; ***, p ≤ 0.001; ns, not significant).

Finally, targeted amino acid analysis confirmed slightly decreased levels of several amino acids in MUT1, although for mostly different individual amino acids (alanine, isoleucine, leucine, proline, threonine, and valine) than the ones found by untargeted metabolomics (only proline was consistently decreased in all analyses), while sarcosine was slightly, but significantly increased in MUT1 samples **(Fig. 6D)**. All other amino acids measured by our targeted method were not significantly changed between wildtype and MUT1 samples (**Supplementary Fig. 9C**). In contrast to the untargeted analysis, two independent targeted methods both showed slightly increased carnitine and acylcarnitine levels in MUT1 versus wildtype samples (**Supplementary Fig. 10**). Given that the targeted and untargeted analyses were performed on samples derived from larvae belonging to different generations, it is difficult to conclude at this stage whether any discrepancies observed between the two approaches have a methodological or biological origin. Many metabolic changes were in agreement between the two methods and those should be considered as the most robust *cln3*-deficiency induced changes that can be detected in zebrafish larvae, in a generation-independent manner. It remains to be established whether acylcarnitine levels are consistently affected by this genetic deficiency in zebrafish as well.

### Further evidence for perturbations in lipid metabolism in *cln3* mutant larvae by lipidomics

We performed HRMS-based untargeted lipidomics analyses on the nonpolar extracts of the discovery and validation sets and detected 11447 MS features (4927 in negative mode and 6520 in positive mode), of which a total of 1901 were annotated based on MS2 spectra (181 in negative and 1720 in positive ionization mode) (**Supplementary Table 4**). We identified 39 lipids with a - fold change >2 and p<0.001. Diverse lipids from different classes (**Supplementary Table 5**), including glycerolipids, were altered in MUT1 larvae **(Fig. 7A, B)**. In particular, a bis(monoacylglycero)phosphate (BMP 22:6_22:6) and acyl steryl glycoside (ASG 27:1;O;Hex;FA 14:1) displayed highly significant changes (>3-fold). To confirm our results, we performed targeted lipidomics analysis of 20 different lipid classes in MUT1 and WT larvae using LC-MS in multiple reaction monitoring mode. This targeted analysis confirmed a global decrease of BMP species at 5 dpf in MUT1 larvae, with the highly abundant BMP 22:6_22:6 showing the most important -fold change **(Fig. 7C)**. Interestingly, while individual cholesterol ester species (CE) changes between MUT1 and WT larvae were not significant, the summed level of cholesterol esters was significantly increased in MUT1 larvae **(Fig. 7D)**. Furthermore, triacylglycerides, hexosyl- and lactosyl-ceramides also showed (non-significant) elevations in MUT1 larvae. In future work, it will be interesting to test whether these early trends for certain lipid classes become more pronounced at later developmental stages. We did not observe any significant changes for ceramides or dihydroceramides, indicating that only the glycosylceramides are affected, which is in line with the acyl steryl glycoside accumulation that was observed in the untargeted analysis. Steryl glycoside formation in animals has been reported to be catalyzed by glucocerebrosidases via transglycosidation of glycosylceramides (Akiyama et al. 2020; Shimamura 2020). Finally, no significant differences were observed for diacylglycerols (DG), plasmologens (PL), sphingomyelins (SM), phosphatidylcholines (PC), phosphatidylethanolamines (PE), phosphatidylglycerols (PG), phosphatidylinositols (PI), phosphatidylserines (PS), lysophosphatidylcholines (LPC), and lysophosphatidylethanolamines (LPE) through the targeted analyses (**Supplementary Fig. 11**).

**Figure 7.**
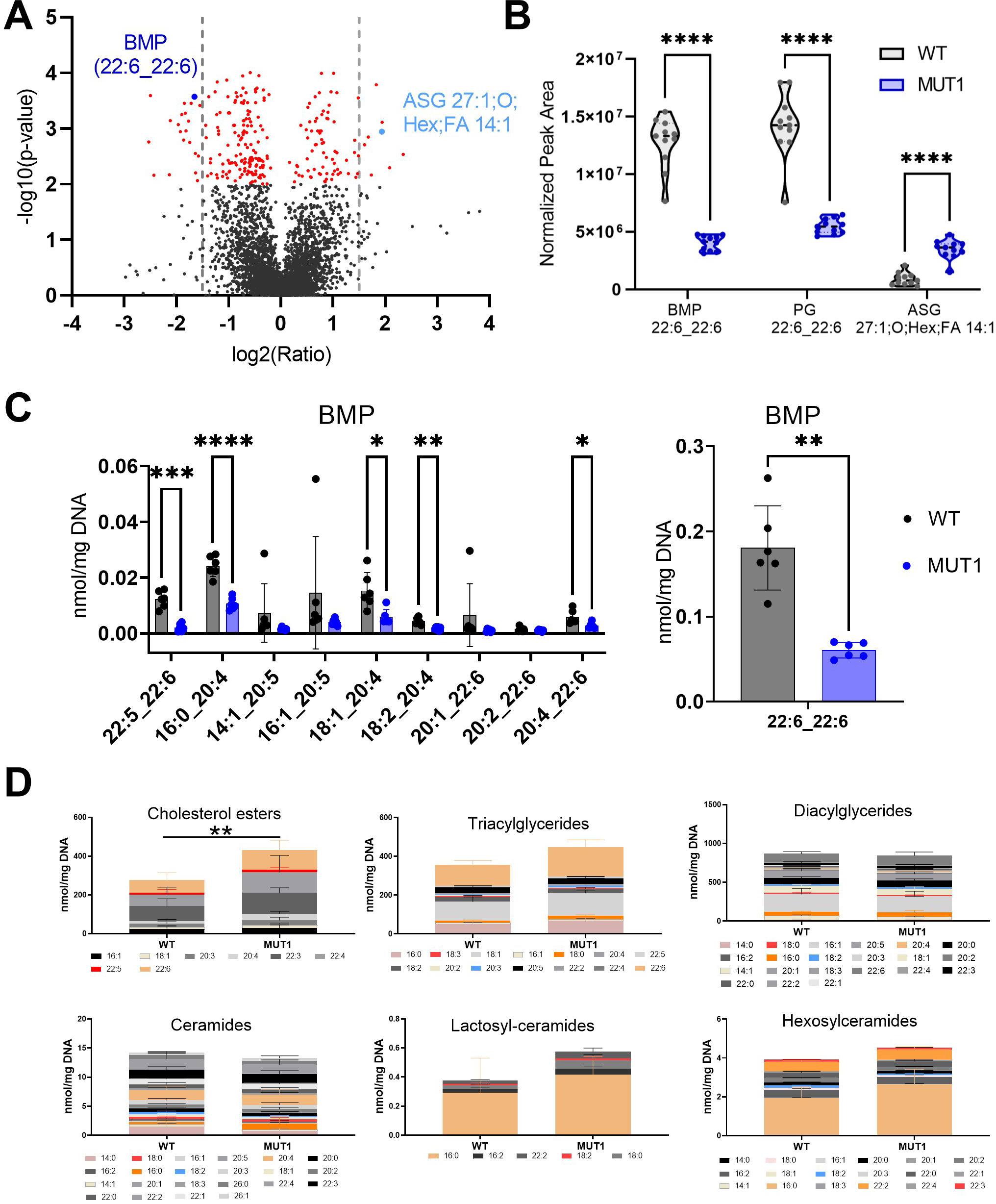
Lipidomics analysis in WT and MUT1 larvae. **(A)** Volcano plot with annotated lipids found by untargeted lipidomics in nonpolar extracts derived from the zebrafish larvae samples generated in the discovery and validation sets as described in the main text. Each dot represents a metabolite and lipids with a significantly different abundance in WT and MUT1 extracts are shown in red (p< 0.001). Grey dotted lines indicate the 1.5 -fold change log2 ratio cut-off. BMP (44:12) and ASG (27:1;O, Hex FA14:1) are highlighted in dark and light blue, respectively. **(B)** Violin plots for the most significantly altered lipids based on untargeted lipidomics analysis. **(C)** Targeted BMP analysis in zebrafish extracts at 5 dpf. Data shown are means ± SDs for 4 biological replicates. Statistically significant differences between WT (grey) and MUT1 (blue) were determined using an unpaired multiple Welch’s t-test (*, p ≤ 0.05; **, p ≤ 0.01; ***, p ≤ 0.001; ****, p ≤ 0.0001). **(D)** Stacked bar graphs for the indicated lipid classes measured by targeted lipidomics in 5 dpf zebrafish larvae. Each stacked bar is the sum of the average amounts measured for the indicated lipid species in six biological replicates of a pool of 40 larvae, normalized against its DNA concentration. Error bars represent SDs. Statistically significant differences between both zebrafish lines were determined using a two-way ANOVA test on the summed averages for each lipid class. **, p ≤ 0.01).

## Discussion

In the present study, we aimed at investigating the function of *CLN3* using zebrafish as *in vivo* model. In a first approach, we knocked down *cln3* using three different morpholino oligonucleotides (one translation-blocking MO and two splice-blocking MOs), but an obvious embryonic phenotype could not be observed in any of the generated morphants after injection of subtoxic MO doses. This contrasted with the severe phenotypes previously published in another study (Wager et al. 2016) in which similar MO target sites on the *cln3* gene were selected, although the exact MO sequences and the genetic background of the zebrafish strains differed. While we used the AB strain for our studies, Wager and colleagues (Wager et al. 2016) used the TLF (Tüpfel Long Fin) strain. The latter is also a commonly used WT line that carries two known mutations, *leo^t1^* and *lof^dt2^*, leading to changes in the pigmentation pattern and the development of long fins, respectively. Although studies in mouse models of *CLN3* disease have highlighted the relevance of the genetic background aspect on phenotype development (Kovacs and Pearce 2015), it seems unlikely that solely background effects could lead to such divergent results as observed here and previously.

The efficiency of our TB-MO could not be validated due to absence of specific antibodies against zebrafish Cln3. However, we could confirm that both of our SB-MOs very efficiently interfered with normal *cln3* transcript generation. As both our translation- and splice-blocking MO treatments remained phenotypically silent, but also to circumvent the ever-lingering doubts about off-target effects surrounding studies that are exclusively MO-based, we generated stable *cln3* mutant lines using the CRISPR-Cas9 technology. In agreement with our morpholino results, both CRISPR homozygous *cln3* mutant lines generated here displayed normal development and reached adulthood without any obvious phenotype. Our observations are also consistent with results obtained in murine models, in which *Cln3* deficiency was not lethal (Kovacs and Pearce 2015). We cannot exclude that the *cln3* transcript variants expressed in the SB-MO-treated morphants as well as in the two generated CRISPR lines may conserve some residual function, as it was reported for the *Cln3* murine models and the most common 1.02 kb deletion variant found in JNCL patients (Kitzmuller et al. 2008). However, the truncated protein predicted to be expressed in the MUT1 line contains only the first transmembrane domain and is much shorter than the predicted expression product of the 1.02 kb deletion variant; therefore, we believe that we have generated a loss-of-function mutation in this MUT1 line. We could not detect a decrease of *cln3* mRNA levels in MUT1, reducing the chance that transcriptional adaptation mechanisms (triggered by mutant mRNA decay) are compensating *cln3* loss-of-function phenotypes in our zebrafish model (El-Brolosy et al. 2019). Finally, the very distinct accumulation of GPDs and decrease of BMP species in MUT1 larvae confirm that we have indeed successfully knocked out Cln3 function, as the exact same metabolic changes were measured in samples derived from patients with CLN3 disease (Hobert and Dawson 2007; Brudvig et al. 2022; Laqtom et al. 2022). The association between BMPs and CLN3 has been known for many years, with metabolic labeling studies showing early on that CLN3 may be involved in BMP synthesis (Hobert and Dawson 2007). By contrast, the accumulation of GPDs in CLN3 deficient mammalian cells and tissues as well as in extracellular fluids (CSF and plasma) of CLN3 patients was only shown very recently in two independent studies (Laqtom et al. 2022; Brudvig et al. 2022), while the work presented here was ongoing in the zebrafish and human cerebral organoids. It is remarkable that we were able to detect GPD accumulation by untargeted metabolomics in an unbiased way (before results from the other groups were published) and in extracts prepared from whole 5 dpf larvae. In the *Cln3* knockout mouse model, tissular GPD accumulation became evident only in samples enriched in lysosomes (lysoIP approach) and was shown for approximately 7-months old animals (Laqtom et al. 2022). In this latter study, evidence is also presented for cultured cells that CLN3 function is required for efflux of GPDs from the lysosomes to the cytosol, where they are normally further catabolized during phospholipid catabolism (Laqtom et al. 2022).

BMP is enriched in the inner leaflet of endo-lysosomal membranes and its principal function remains unclear. Also, little is known about the pathways involved in BMP formation and degradation, although its synthesis is predicted to occur in the late endosomes/endo-lysosomes, as its concentration increases over the maturation of the latter (Gruenberg 2020). BMP is highly enriched in intraluminal vesicles (ILVs) of multivesicular bodies (MVBs), which are involved in cholesterol efflux and sphingolipid degradation (Gruenberg 2020). Accordingly, BMP seems to play a key role in lipid metabolism and membrane composition by facilitating the adhesion of soluble, positively charged hydrolases and activator proteins such as sphingolipid activator proteins SapsA-D and the GM2 activator protein to intracellular membranes (Gallala and Sandhoff 2011). It is therefore possible that the BMP depletion in CLN3 deficient cells also negatively affects the function of glucocerebrosidases, which may then possibly explain the increased levels of glycosyl-ceramide and acyl steryl glycoside species that we observed in MUT1 larvae (Ben Bdira et al. 2018). Moreover, BMPs have a cone-like shape that affects membrane curvature, playing an important role for endosomal vesicularisation and hence also in the maturation of late endosomes (Gallala and Sandhoff 2011). In summary, CLN3 deficiency leads to profound changes in the lipidome, particularly in lipids associated with the endo-lysosomal compartment (BMP depletion and GPD accumulation), already during early development, which may compromise endo-lysosomal membrane integrity and transport mechanisms across these membranes, as well as cause further perturbations in endo-lysosomal metabolic activities, likely to lead progressively to dysfunctional lysosomes accumulating also other lipids (e.g. cholesterol esters, triglycerides, glycosyl- and lactosyl-ceramides) to abnormal levels **(Fig. 8)**.

**Figure 8.**
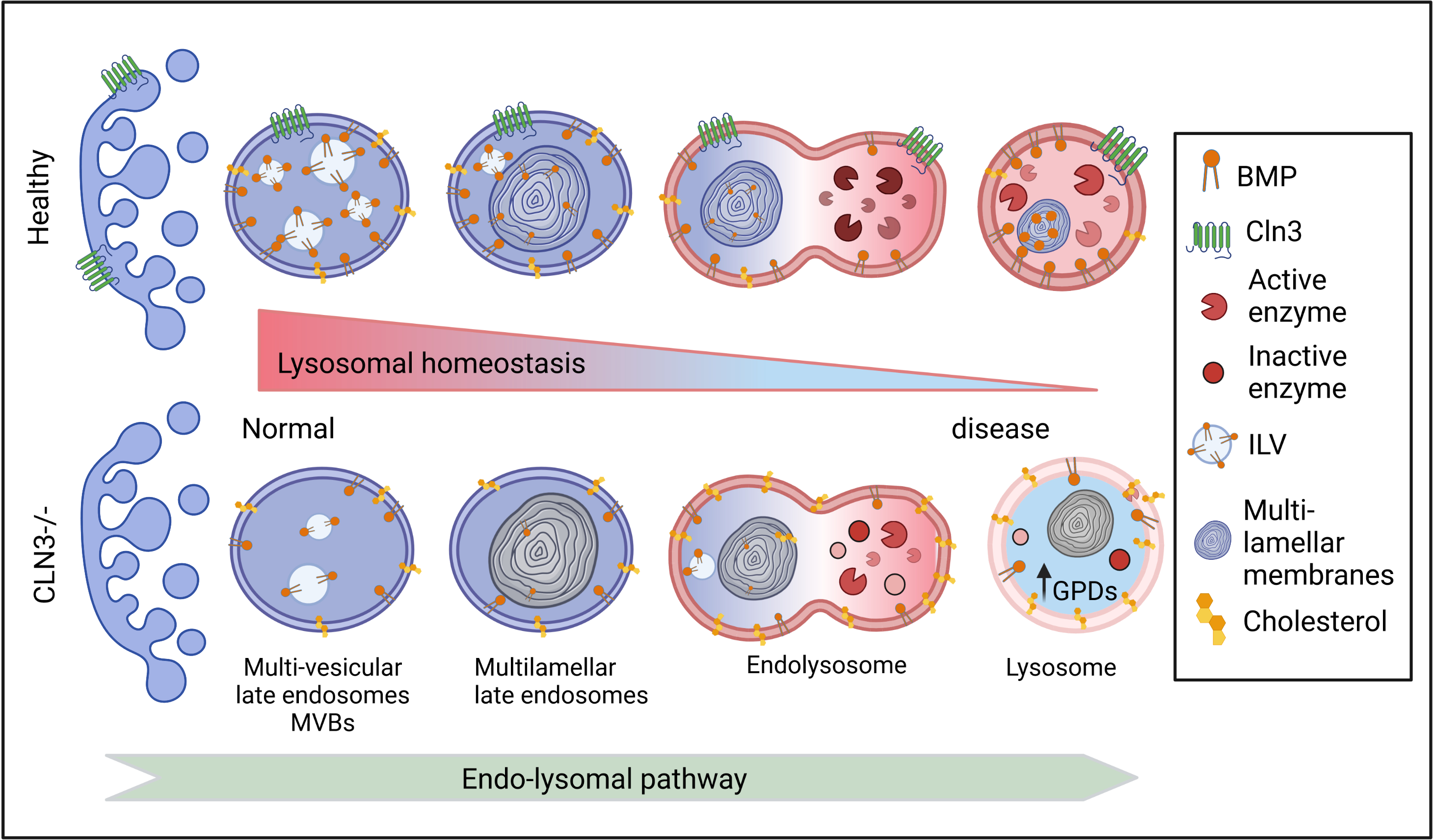
Proposed changes induced by CLN3 deficiency along the endo-lysosomal pathway. In healthy cells, the number of intraluminal vesicles (ILVs) increases in the early endosomes to form multilamellar bodies in the late endosomes. The latter fuse with lysosomes forming a hybrid transient endo-lysosome to finally form secondary lysosomes. BMP is enriched in ILVs and its levels increase during endosome maturation, contributing to ILV formation, ceramide degradation, and cholesterol efflux (Gruenberg 2020). In CLN3 deficient cells, BMP levels are decreased, potentially leading to reduced ILV formation as well as impaired degradation and therefore accumulation of cholesterol, glycosylceramides and glycerolphosphodiesters (GPDs), all contributing to lysosomal dysfunction.

Although the metabolic perturbations that we found in our *cln3* mutant zebrafish larvae are perfectly in line with findings in samples derived from patients with CLN3 disease, the behavioral phenotypes observed in these larvae were surprisingly subtle when compared to the severe neurological symptoms that are, unfortunately, commonly observed in the patients. However, the changes that we could find were robust and, given that we observe them very early in development, may help shed light on critical neurodevelopmental processes that are impaired in CLN3 patients, well before the symptoms manifest. Locomotor behavioral assays at 5 dpf showed that the MUT1 larvae have higher basal activity, in both light and dark conditions, compared to control larvae. In contrast, the MUT2 line showed only (non-significantly) increased activity in the light condition. In the dark-light response assay, both homozygous mutant lines showed a much less pronounced startle response upon the lightening change, possibly indicating a visual acuity defect in the mutants. Considering that visual impairment is the first obvious symptom to manifest in juvenile BD patients (Ostergaard 2016), it will be very important to clarify whether this phenotype has a retinal or neuronal origin.

Our metabolomics analyses identified dysregulation in the levels of three important neurotransmitters, NAAG, glutamine and GABA, further suggesting that brain homeostatic functions are perturbed in *cln3* mutant zebrafish. Attenuation of NAAG levels in the MUT1 line corroborates previous results obtained with CSF from *CLN2* patients and animal models for Batten Disease (Sindelar et al. 2018; Elshatory et al. 2003; Gomez-Giro et al. 2019; Pears et al. 2005). Neurotransmitter dysregulation might be responsible for epileptic seizures observed in 50% of the children with *CLN3* disease (Schulz et al. 2013; Bosch and Kielian 2019). Studies in *Cln3^lacZ/lacZ^*mice revealed that they are more prone to develop seizures when treated with subacute doses of the chemoconvulsant PTZ (Eliason et al. 2007). We tested PTZ in 5 dpf zebrafish larvae, and we observed a significant increase in locomotor activity in the MUT1 line compared to control larvae at subacute concentrations. Interestingly, an even more pronounced phenotype was observed upon exposure to PTX. Our results suggest that *cln3* mutants are more sensitive to PTX, since MUT1 larvae showed hyperactivity already at 0.1 mM, whereas in control larvae the hyperactivity only occurred starting at 0.3 mM. Strikingly, at PTX concentrations higher than 0.3 mM, the mutant activity significantly dropped compared to the wildtype larvae. As described for PTZ (Baraban et al. 2005), wildtype behavior under exposure to PTX can be divided in three stages. In the first minutes, the swimming activity increases (stage I, ≈ 0-10 min), followed by a rapid “whirlpool-like” circling swimming behavior (stage II, ≈ 10-26 min), and finally a progressive decrease of the activity (Stage III, ≈ 26-60 min). The latter stage is characterized by clonus-like convulsions and loss of posture that unfortunately cannot be accurately detected by the automated behavioral tracking system that we used. However, the decrease in movement that we observed in our experiments certainly suggests that the larvae had entered stage III. Strikingly, both *cln3* mutants clearly showed stages I and III, but stage II seemed to be very short or completely absent. One interpretation is that the *cln3* mutant larvae skip most of the stage II reaction (“whirlpool-like” circling swimming) and directly enter stage III, in which the larvae undergo repeated episodes of convulsions (Baraban et al. 2005). In this stage, the larvae display clonus-like seizures reflected by rapid movement followed by postural loss where larvae turn upside down and remain immobile (Baraban et al. 2005). The skipping of stage II would explain the decrease of the total movement that we observed with both *cln3* mutant lines.

No obvious abnormalities were observed in the MUT1 and MUT2 lines at adult stage, which may be explained by the remarkable ability of zebrafish to regenerate neuronal cells (Alunni and Bally-Cuif 2016; Kizil et al. 2012) and/or the induction of other compensatory mechanisms. In contrast to mammals, the zebrafish adult brain is enriched with a wide variety of neurogenic niches that constantly generate new neurons (Zambusi and Ninkovic 2020). Moreover, inflammation caused by brain injury or by induced neurodegeneration activates specific neurogenic programs that efficiently overcome the loss of neurons, allowing the maintenance of the tissue architecture in zebrafish brain (Diotel et al. 2020). This regenerative potential of zebrafish may mask more pronounced loss-of-function phenotypes expected to develop with increasing age in *CLN3* deficiency.

Altogether, our MUT1 larvae recapitulate, within the first 5 days of life, metabolic abnormalities observed at a more advanced age in other *CLN3* models and patients, provide further support for perturbed lipid metabolism in the pathogenesis of *CLN3* disease, and highlight BMPs and GPDs as promising biomarker candidates for very early detection of this disease, maybe long before symptoms manifest. Our findings at larval stages open new avenues for investigation of the physiological function of CLN3 in an *in vivo* model that is more time- and cost-efficient than other whole organism models. The observation that GPDs accumulate to high levels in *CLN3* deficient zebrafish larvae and human cerebral organoids, *i.e.* during early developmental stages, suggests that GPD metabolism is very tightly connected to CLN3 function. The absence of GPD accumulation in our zebrafish model for ATP13A2/CLN12 deficiency (Heins-Marroquin et al. 2019), also leading to neuronal lipofuscinosis (Farias et al. 2011; Bras et al. 2012; Matsui et al. 2013), indicates that these compounds may be specific for certain types of NCLs, a hypothesis that will now need to be tested in different models and non-CLN3 Batten disease patients. The zebrafish model has the potential to fill the gaps between cell culture and mammalian models and accelerate the understanding of early metabolic events that lead to the development of a childhood disease that, as of today, is still invariably lethal.

## Materials and methods

### Ethics statement

The Zebrafish Facility at the Luxembourg Centre for Systems Biomedicine is registered as an authorized breeder, supplier, and user of zebrafish with Grand-Ducal decree of 20 January 2016 and 26 January 2023. All practices involving zebrafish were performed in accordance with European laws, guidelines and policies for animal experimentation, housing and care (European Directive 2010/63/EU on the protection of animals used for scientific purposes). Authorization number LUPA 2019/01 allowed the generation of the *cln3* mutant lines in this project, and authorization number LUPA 2017/04 allowed the performance of fin biopsies for genotyping purposes. Experiments using larvae older than 5 dpf were performed under Grand-Ducal decree of 10 December 2012.

### Zebrafish lines and handling

Wildtype zebrafish (AB) and mutant lines (*nacre*, *cln3^lux1^*, *cln3^lux2^*, *atp13a2^sa18624^*) were kept in the Aquatic Facility of the Luxembourg Centre for Systems Biomedicine (LCSB) according to standard protocols (Westerfield 2000). Zebrafish embryos were obtained by natural spawning and reared in 0.3X Danieau’s solution (17 mM NaCl, 2 mM KCl, 0.12 mM MgSO_4_, 1.8 mM Ca(NO_3_)_2_, 1.5 mM HEPES pH 7.5, and 1.2 µM methylene blue) in 10h dark – 14h light conditions at 28°C. Adult fish were maintained in 3.5 l tanks at a stock density of 4-6 fish/l with the following parameters: water temperature, 28.5 °C (±0.5); light:dark cycle, 14h:10h; pH 7.5; conductivity, 700-900 µS/cm. Fish were fed three to five times a day, depending on age, with granular and live feed (*Artemia salina*). For live imaging, larvae were first anesthetized with buffered 0.008% tricaine methane sulfonate (MS222), and then embedded in 2.5% methylcellulose.

### Morpholino antisense oligonucleotide microinjection

Translation blocking (TB) and splice blocking (SB) morpholinos (Mos) for *cln3* were purchased from Gene Tools (Oregon, USA) with the following sequences: CLN3_SB-e2i2, 5’-GAT TGA CTC AAC ATT ACC AGA ACG C-3’; CLN3_TB-ATG, 5’-TCG ATC CAT TGC GAC TTT CAC AGG A-3’; and CLN3_SB-i2e3 5’-CCC CAG CAA CCT AAA CAG AGA TAA T-3’. A standard MO with the sequence 5’-CCT CTT ACC TCA GTT ACA ATT TAT A-3’ was used as a non-targeting MO control. All morpholinos had a 3’-lissamine or -fluorescein modification. Stock solutions (2 mM) were prepared according to the specifications of the provider and titrated working solutions were freshly prepared for each experiment. Manual microinjections of Mos were performed as previously described (Rosen, Sweeney, and Mably 2009). Briefly, gene knockdown was achieved through microinjection of 1 nl of *cln3*-MO or control-MO into the yolk of wildtype embryos at one- to two-cell stage using an Eppendorf FemtoJet 4X® microinjector. Injected embryos were incubated at 28°C, and fluorescently sorted and evaluated daily until needed for the analytical experiments.

### CLN3 sequence alignment

Multiple sequence alignment was performed with the MUSCLE software from EMBL-EBI using the following CLN3 protein sequences: *Saccharomyces cerevisiae* (NP_012476.1), *Danio rerio* (NP_001007307), *Homo sapiens* (NP_001273038), *Mus musculus* (NP_001139783). The Jalview software was used for editing the alignment for visualization purposes.

### Generation of CRISPR/Cas9 mutants

#### sgRNA template generation and transcription

The target sequences in the *cln3* and *slc45a2* genes were selected using the online tool CHOPCHOP (http://chopchop.cbu.uib.no/index.php). For generation of the sgRNA templates for *cln3* and *slc45a2*, 60-nt gene specific oligos were designed. Both oligos contained a T7 (5’-TAATACGACTCACTATA-3′) promoter sequence, a 20-nt target site without the Protospacer Adjacent Motif (PAM), and a 23-nt complementary region to the tracrRNA tail. The oligos were annealed to a generic oligonucleotide encoding the reverse-complement of the tracrRNA tail. Subsequently, the ssDNA overhangs were filled in with T4 DNA polymerase (NEB) following the manufacturer’s instructions. The resulting templates were purified using the Wizard® SV Gel and PCR Clean-Up System kits (Promega) and transcribed using the Megascript T7 kit (Thermo Scientific) according to the manufacturer’s instructions. The transcription reaction was performed for 4 hours and the preparation was then treated with 1 µl Turbo Dnase for 15 min at room temperature. The sgRNAs were precipitated with ammonium acetate, resuspended in 50 µl water, and the concentration was determined by measuring A_260_ (NanoDrop 2000, Thermo Scientific). All nucleotide and primer sequences used in this study are listed in **Supplementary Table 6**.

#### sgRNA and Cas9 protein microinjection

200 ng of *cln3* sgRNA (0.5 μl), 200 ng of *slc45a* sgRNA (0.5 μl), 160 ng Cas9 Nuclease NLS from (A) *S. pyogenes* (1 μl, NEB), and 0.2 µl phenol red were mixed in an Eppendorf tube (total volume 2.2 μl) and incubated for 5 minutes at room temperature. The mixture was placed on ice and 1 nl was injected into one-cell embryos (Rosen, Sweeney, and Mably 2009). *Slc45a2* sgRNA was used to track successful injections with albino clones indicating that Cas9 had worked efficiently. Chimeras showing an albino phenotype were selected and raised in the LCSB Aquatic Facility as described above.

#### PCR analysis from whole larvae

Larval genomic DNA was prepared using the HotSHOT protocol (Meeker et al. 2007). Briefly, single injected 1 dpf embryos were dissociated in PCR tubes containing 10 µ1 of 50 mM NaOH at 95 °C for 20 min. After cooling down, 1 µl Tris-HCl pH 7.5 was added and the preparation was mixed. 1 µl of this mixture was used as genomic DNA template in PCR reactions using the primers listed in S6 Table. Amplicons were analyzed on 2.5% agarose gels prepared with 1× Tris-acetate-EDTA buffer (Sigma).

#### PCR analysis from adult fish by fin biopsy

Adult zebrafish were anesthetized in 0.016% MS222. A small portion of the caudal fin was cut, placed in an Eppendorf tube containing 250 µl lysis buffer (100 mM Tris-HCl pH 8, 200 mM NaCl, 0.2% SDS, 5 mM EDTA pH 8) supplemented with 0.1 mg/ml proteinase K, and digested at 55°C for 2 h or overnight with vigorous shaking. Proteinase K was inactivated for 20 min at 90°C and 400 µl of isopropanol was added to the cooled samples, followed by vortexing and centrifugation at 13000 *g* and 4°C for 30 min. The pellet was washed with 100 µl of 70% ethanol and then dried at 55°C for 10 min. The washed pellet was resuspended in 50 µl water and finally incubated 10 min at 55°C to dissolve the DNA. The nucleic acid concentration was determined by measuring A_260_ (NanoDrop 2000, Thermo Scientific) and 100 ng DNA were used per PCR reaction (final volume 30 µl). The PCR products were analyzed by electrophoresis on a 2.5% agarose gel. For mutation identification, PCR amplicons were purified using the Wizard® SV Gel and PCR Clean-Up System kits (Promega), followed by Sanger sequencing (Eurofins). The DNA sequences were compared to the wildtype *cln3* sequence (ENSDART00000055170) listed in the ENSEMBL database. Primer sequences are given S6 Table.

### cDNA sequencing

Total RNA extraction and cDNA synthesis were performed as described previously (Heins-Marroquin et al. 2019). 1 μl cDNA of WT and CRISPR *cln3* mutant lines was used as PCR template and amplicons were cloned into the pGEMT vector following the manufacturer’s instructions (Promega). Selected clones were sent for sequencing.

### Locomotor behavior assays

Locomotor activity of 5 dpf larvae in response to light-dark conditions or pro-convulsion treatment was analyzed using a DanioVision system equipped with a temperature control unit (Noldus Information Technology). The EthoVision XT 11.5 software (Noldus Information Technology) was used to monitor the movements of individually swimming larvae. Motion values were expressed as mean swimming velocity (mm/s). All tracking experiments were performed at least as independent triplicates (with at least 8 larvae per genotype per replicate). Technical outliers (*i.e.* no detection of the larva by the system) were removed from the quantification. Statistical analysis was performed using the GraphPad Prism analysis software (version 8.2.0).

#### Dark-light-dark and global activity tracking

One hour prior to the recording, 5 dpf zebrafish larvae were carefully transferred into each well of a 24-well plate, with 1 ml Danieau’s medium per well. The plate was then incubated at 28°C in the dark. For activity recording, each plate was placed in the DanioVision system and the larvae were allowed to habituate in the dark to the new environment for 15 min. After this period, locomotor activity was continuously tracked using a dark-light-dark protocol (15-min intervals). For the global activity test, larvae were placed in the DanioVision system to habituate in the tested lightening condition for 15 min, followed by locomotor activity recording for 45 min. All tested genotypes were represented by the same number of larvae in each 24-well plate to avoid any variations due to differences in experiment timing and handling. All experiments were performed during the day between 2 and 6 pm.

#### Pro-convulsant treatment

For each experimental group, twelve 5 dpf zebrafish larvae were placed individually in 96-well plates. Each well contained 100 μl of Danieau’s medium. Subsequently, 100 μl of pro-convulsant working solution (PTX or PTZ) was added to each well to obtain final concentration ranges of 0.01-1 mM for PTX and 5-20 mM for PTZ. Immediately after addition of the drug or 10 min later, locomotor behavior was recorded in the DanioVision system for 1h in the dark.

### RNA extraction and qPCR

Adult zebrafish were euthanized in buffered 0,04% MS222 before organ dissection and 30 larvae at the indicated developmental stages were euthanized by hypothermia. For total RNA extraction, organs and larvae were placed in Precellys tubes containing 1.4 mm ceramic beads and 900 μl TRIzol (Thermo Fisher). Samples were homogenized for 30 sec at 6000 rpm (Precellys24, Bertin Instruments) and at 4°C and 200 µl chloroform was added to the lysates, followed by centrifugation for 15 min at 15700 *g* and 4 °C. The upper aqueous phase was transferred into a fresh Eppendorf tube containing 450 μl isopropanol, followed by vortexing. After a 10-min incubation at room temperature, samples were centrifuged again for 15 min at 13000 *g* and 4 °C. Pellets were washed with 1 ml 70% ethanol and dried for 5-10 min. Finally, pellets were resuspended in 30 µl MilliQ water, the RNA concentration determined by measuring A_260_ (NanoDrop 2000), and the RNA quality evaluated by agarose gel electrophoresis. After treatment with Dnase I (Rnase free, Sigma), total RNA (2 µg per reaction) was reverse transcribed with the SuperScript III reverse transcriptase (Invitrogen) according to the manufacturer’s instructions. Ten-fold diluted cDNA samples (2 μl) were added to a mixture (10 μl total volume) containing iQ™ SYBR® Green Supermix (5 μl; Bio-Rad) and the primers of interest at a final concentration of 10 μM. Gene expression was analyzed by quantitative PCR (qPCR) in 384-well plates using the LightCycler480 (Roche). For each tissue, qPCR was performed for three biological replicates, with two technical replicates per biological replicate. Expression levels of *cln3* in the indicated tissues were calculated relative to the eye tissue using the 2^ᐃᐃCt^ method (Livak and Schmittgen 2001). *Ef1α* and *rpl13α* were used as reference genes for normalization. All qPCR primer sequences are given in S6 Table.

### Untargeted metabolomics analysis using HILIC-HRMS

Zebrafish larvae at 5 dpf were washed twice with 2 ml 0.3X Danieau’s solution and transferred to labeled and pre-weighed Precellys tubes (40 larvae/tube). Larvae were euthanized on ice and stored at -80 °C. Before extraction, samples were lyophilized overnight then weighed. Cold dichloromethane (DCM, 330 µl), methanol (170 µl) and 25 ceramic beads (ᴓ 1.4 mm) were added to each tube and samples were homogenized twice (3000 rpm for 30s) at -10 °C using a Precellys instrument (Bertin Instruments). After vortexing, homogenates (400 µl) were transferred to 2-ml Eppendorf tubes containing 117 µl of cooled Milli-Q water, followed by mixing through inversion. In a thermomixer, samples were further mixed for 20 min at 1000 rpm and 4 °C. Finally, samples were centrifuged at maximum speed (16000 *g*) for 20 min at 4 °C and 100 µl of the upper aqueous layer were collected for metabolomics analysis whereas 100 µl the organic lower phase were collected for lipidomics analysis. The non-polar extracts were vacuum concentrated until dryness (CentriVap Benchtop Vacuum Concentrator, LABCONCO).

Hydrophilic interaction chromatography (HILIC) was carried out using a Sequant Zic-pHILIC column (5 μm, 2.1 × 150 mm) equipped with a guard column of the same chemistry (5 μm, 2.1 × 20 mm). The flow rate was set at 0.20 ml/min using aqueous 20 mM ammonium acetate, pH 9.0 (A) and 20 mM ammonium acetate, pH 9.0 in acetonitrile (B) as mobile phases. The mobile phase gradient started at 90% B for 1.5 min before linearly ramping to 20% B over 14.5 min. This condition was maintained for 2 min before returning to the starting mobile phase conditions over 2 min. The column was allowed to re-equilibrate for 10 min at 90% B before the next injection. Mass spectrometry was performed separately in positive and negative mode using a High Field Q Exactive mass spectrometer (Thermo Scientific). MS1 data acquisition settings were as follows: mass resolution of 120000, default charge 1, automatic gain control (AGC) 1×10^6^ with maximum injection time (IT) of 70 ms over a scan range of 60-900 *m/z*. For MS/MS acquisition, MS/MS events were triggered in data-dependent dynamic exclusion fashion where the top 5 highest intensity ions were fragmented with a dynamic exclusion window of 10 s. The AGC was set to 8×10^3^ with a maximum IT of 70 ms. Minimum ion intensity was 1×10^5^ with an isolation window of 1 Da and HCD collision energy of 30 NCE (nominal collision energy).

Untargeted data analysis was performed using Progenesis QI (version 2.2; Waters) for the two different ionization modes separately. For peak picking, a minimum peak width of 0.1 min was used. Three consecutive metabolite annotation searches were performed with the Progenesis software. The first round used the provided METLIN MS/MS database, while the second round used the following subset of selected databases for MS1 and (*in silico*) MS2 annotations within the ChemSpider mode: BioCyc, Human Metabolome DataBase (HMDB), Kyoto Encyclopedia of Genes and Genomes (KEGG), LipidMaps, PubMed, Peptides and Yeast Metabolome DataBase (YMDB). In the last round, a LipidBlast search was performed (Kind et al. 2013). Only proton loss ([M-H]^−^) and proton gain ([M+H]^+^) were considered during peak annotation for negative and positive modes, respectively. The mass error for MS1 and MS2 searches were both set at 5 ppm. For the remaining Progenesis settings, the default setting was selected. Sample normalization was performed using ‘the normalize to all compounds’ method provided by the Progenesis software. Quantitative extracted ion chromatogram areas together with the primary metabolite annotations for both the discovery and validation batches are available in S1 and S2 Tables, respectively. The annotation with highest Progenesis score was chosen as primary annotation. In case of identical Progenesis scores, the alphabetically first annotation was designated as the primary annotation. All (redundant) compound annotations for both batches together with Progenesis scores and MS2 identification scores (when available) are given in **Supplementary Table 7** together with ChemSpider or METLIN identifiers. Selected metabolites of interest were validated with standard compounds (S3 Table).

Univariate and multi-variate statistical analyses were performed using the statistical program R (ver. 4.01). For univariate analysis, the extracted ion chromatogram areas for the annotated features were log10 transformed and t-tested using the packages *ClassComparison* (ver 3.1.8) and *stats* (ver 4.0.2). For multivariate analysis, principal component analysis (PCA) was performed using the *FactoMineR* package, ver 2.3 (Galili et al. 2018). Metabolite enrichment analysis was performed using the *MetaboAnalystR* R-package (ver 3.0.2) (Pang et al. 2020) including statistically significant differential features that had a primary annotation that could be mapped to an HMDB identifier.

### Targeted metabolomics analyses using LC-MS

Deuterated amino acid and carnitine mixture (NGS Internal Standard, 57004), used as internal standard, was reconstituted in 80% methanol and further diluted according to the NGS Internal Standard 57004 manual. Extraction fluid was prepared by adding four parts methanol to one part NGS Internal Standard stock solution. For metabolite extraction, 1 mL extraction fluid was added to 40 zebrafish larvae and homogenized using a Precellys Evolution homogenizer (Bertin Technologies) in the presence of 500 mg ceramic beads (1.4 mm) for 30 sec at 6000 rpm and 0 to 5 °C. The homogenate was incubated at 4 °C with shaking (1400 rpm) for 10 min (Eppendorf Thermomixer), followed by centrifugation at 21,000×g for 10 min at 4 °C. Then, 70 µL of the supernatant were filtered and transferred into LC vials with micro insert for amino acid and glycerophosphodiester analysis. In addition, 100 µL of the supernatant were dried in a rotary vacuum concentrator at -4 °C. Dried extracts were reconstituted in 100 µL water/acetonitrile (1:1, v/v), filtered and submitted to carnitine analysis.

For amino acid and glycerophosphodiester analysis, measurements were performed using a UHPLC Vanquish system coupled to an Orbitrap Exploris 240 mass spectrometer (ThermoFisher Scientific) equipped with an OptaMax NG electrospray ionization source. Chromatography was carried out with a SeQuant ZIC-pHILIC 5µm polymer (150 × 2.1 mm) column protected by a SeQuant ZIC-pHILIC Guard (20 × 2.1 mm) pre-column. The column temperature was maintained at 45 °C. Mobile phases consisted of 20 mmol/L ammonium carbonate in water, pH 9.2, containing 0.1% InfinityLab deactivator additive (Eluent A) and 100% Acetonitrile (Eluent B). The gradient was: 0 min, 80% B; 3 min, 80% B; 18 min, 20% B; 19 min 20% B; 20 min 80% B; 30 min 80% 1. B. The flow rate was set to 0.2 mL/min and 0.4 mL/min for the re-equilibration phase. The injection volume was 5 µL. MS analysis was performed using electrospray ionization with polarity switching enabled (+ESI/-ESI). The source parameters were applied as follows: sheath gas flow rate, 35; aux gas flow rate, 7; sweep gas flow rate, 0; RF lens 70%; ion transfer tube temperature, 320 °C and vaporizer temperature, 275 °C. Spray voltage was set to 3000 V in positive and -3000 V in negative mode. The Orbitrap mass analyser was operated at a resolving power of 60,000 in full-scan mode (scan range: m/z 75-1000; standard automatic gain control; maximum injection time: 100 ms). Data were acquired with Thermo Xcalibur software (Version 4.5.474.0) and analysed with TraceFinder (Version 5.1). Target compounds were identified by retention time, accurate mass and isotopic pattern based on authentic standards. In addition, the identity of all targets was confirmed by accurate mass, MS/MS experiments and standard addition experiments, when authentic standards were available. Briefly, annotation of the compounds of interest was achieved by isolating the precursor ion and fragmenting it into the product ions. The resulting product ion spectrum was used for library matching and qualitative MS interpretation. The procedural error was corrected by using the response ratio of the integrated peak area of the target compound and the integrated peak area of the dedicated internal standard.

For carnitine analysis, measurements were performed using an Exion LC coupled to a 7500 Triple quad MS (SCIEX) equipped with an Optiflow Pro Ion Source. The ion source was operated in electrospray ionization mode. Chromatography was accomplished using a Waters ACQUITY UPLC CSH C18 1.7 µm (2.1 mm × 100 mm) column protected by a VanGuard pre-column (2.1mm × 5 mm). The column temperature was maintained at 40 °C. The mobile phases consisted of 10 mmol/L ammonium formate in water, pH unadjusted (Eluent A) and acetonitrile containing 0.1% formic acid (Eluent B). The flow rate was set to 0.3 mL/min. The LC method consisted of 2-min isocratic delivery of 5% B, a 10-min linear gradient to 95% B and a 3-min isocratic delivery of 95% B, followed by a re-equilibration phase on starting conditions at 5% B for 5 min. The injection volume was 5 µL. Target compounds were measured in scheduled multiple reaction monitoring mode. Specific transitions of each target analyte are given in **Supplementary Table 8**. The source and gas parameters applied were as follows: Ion source gas 1 and 2 were maintained at 40 psi and 70 psi, respectively; curtain gas at 40 psi; CAD gas at 10; source temperature was held at 550 °C. Spray voltage was set to 2000 V in positive ion mode. Mass spectrometric data were acquired with SCIEX OS (Version 3.0.0) and analyzed with MultiQuant (Version 3.0.3). Target compounds were identified by retention time and ion ratio. In addition, the identity of all targets was confirmed by MS/MS. The data set was normalized by using the response ratio of the integrated peak area of target compounds and the integrated peak area of the dedicated internal standard.

### Untargeted lipidomics analysis using LC-HRMS

The non-polar extracts were reconstituted in 100 µl methanol:toluene (9:1). Untargeted lipidomics analysis was adapted from (Cajka and Fiehn 2014). Briefly, lipid separation was accomplished by chromatography on an Acquity UPLC CSH C18 column (1.7 μm, 100 × 2.1 mm) equipped with an Acquity UPLC CSH C18 VanGuard precolumn (1.7 μm, 5 × 2.1 mm) (Waters, Milford, MA) using the same LC-HRMS instrument as for untargeted metabolomics analysis. Column temperature was set at 65 °C and mobile phase flow rate at 0.6 mL/min. Mobile phases (A) 60:40 (v/v) acetonitrile:water with ammonium formate (10 mM) and formic acid (0.1%) and (B) 90:10 (v/v) isopropanol:acetonitrile with ammonium formate (10 mM) and formic acid (0.1%) were mixed according to the following gradient program: 0 min, 15% (B); 0−2 min, 30% (B); 2−2.5 min, 48% (B); 2.5−11 min, 82% (B); 11−11.5 min, 99% (B); 11.5−12 min, 99% (B); 12−12.1 min, 15% (B); and 12.1−15 min, 15% (B). Sample temperature was maintained at 4 °C. Peak picking, lipid MS2 annotation, and data normalization by LOESS algorithm were performed following the protocol for lipid analysis using MS-DIAL (version 4.48) (Tsugawa et al. 2015; Tsugawa et al. 2020).

### Targeted lipidomics analysis using LC-MS

Lipid extraction and targeted lipidomics analysis were performed by Lipometrix at the KU Leuven. An amount of cells containing 10 μg of DNA was homogenized in 700 μL of water with a handheld sonicator and was mixed with 800 μl HCl(1M):CH3OH 1:8 (v/v), 900 μl CHCl3, 200 μg/ml of the antioxidant 2,6-di-tert-butyl-4-methylphenol (BHT; Sigma Aldrich), 3 μl of UltimateSPLASH™ ONE internal standard mix (#330820, Avanti Polar Lipids), and 3 μl of a 10 μg/mL solution of oleoyl-L-carnitine-d3 (#26578 Cayman Chem). After vortexing and centrifugation, the lower organic fraction was collected and evaporated using a Savant Speedvac spd111v (Thermo Fisher Scientific) at room temperature and the remaining lipid pellet was stored at – 20°C under argon.

Just before mass spectrometry analysis, lipid pellets were reconstituted in 100% ethanol. Lipid species were analyzed by liquid chromatography electrospray ionization tandem mass spectrometry (LC-ESI/MS/MS) on a Nexera X2 UHPLC system (Shimadzu) coupled with hybrid triple quadrupole/linear ion trap mass spectrometer (6500+ QTRAP system; AB SCIEX). For acylcarnitine, PG and BMP measurement, chromatographic separation was performed on a Luna® NH_2_ column (100 mm × 2 mm, 3 μm; Phenomenex) maintained at 35°C using mobile phase A [2 mM ammonium acetate in dichloromethane/acetonitrile 7:93 (v/v)] and mobile phase B [2 mM ammonium acetate in water-acetonitrile 50:50 (v/v)] in the following gradient: 0-2 min: 0% B; 2-11 min: 0% B → 50% B; 11-12.5 min: 50% B → 100% B; 12.5-15 min: 100% B; 15-15.1 min: 100% B → 0% B; 15.1-17 min: 0% B, at a flow rate of 0.2 mL/min, which was increased to 0.7 mL/min from 2 to 15 minutes. For all other lipids, chromatographic separation was performed on an Xbridge amide column (150 mm × 4.6 mm, 3.5 μm; Waters) maintained at 35°C using mobile phase A [1 mM ammonium acetate in water-acetonitrile 5:95 (v/v)] and mobile phase B [1 mM ammonium acetate in water-acetonitrile 50:50 (v/v)] in the following gradient: 0-6 min: 0% B → 6% B; 6-10 min: 6% B → 25% B; 10-11 min: 25% B → 98% B; 11-13 min: 98% B → 100% B; 13-19 min: 100% B; 19-24 min: 0% B, at a flow rate of 0.7 mL/min, which was increased to 1.5 mL/min from 13 minutes onwards. Sphingomyelins (SM), cholesterol esters (CE), ceramides (CER), dihydroceramides (DCER), hexosylceramides (HCER), lactosylceramides (LCER) and acylcarnitines were measured in positive ion mode with a production ion of 184.1, 369.4, 264.4, 266.4, 264.4, 264.4 and 85.0, respectively. Triacylglycerides (TAG) and diacylglycerides (DAG) were measured in positive ion mode with a neutral loss for one of the fatty acyl moieties. Phosphatidylcholine (PC), lysophosphatidylcholine (LPC), Phosphatidylethanolamine (PE), lysophosphatidylethanolamine (LPE), phosphatidylglycerol (PG), BMP, phosphatidylinositol (PI) and phosphatidylserine (PS) were measured in negative ion mode by fatty acyl fragment ions. Lipid quantification was performed by scheduled multiple reactions monitoring (MRM), the transitions being based on the neutral losses or the typical product ions as described above. The instrument parameters were as follows: Curtain Gas, 35 psi; Collision Gas, 8 a.u. (medium); IonSpray Voltage, 5500 V and −4500 V; Temperature, 550°C; Ion Source Gas 1, 50 psi; Ion Source Gas 2, 60 psi; Declustering Potential, 60 V and −80 V; Entrance Potential, 10 V and −10 V; Collision Cell Exit Potential, 15 V and −15 V. The following fatty acyl moieties were taken into account for the lipidomic analysis: 14:0, 14:1, 16:0, 16:1, 16:2, 18:0, 18:1, 18:2, 18:3, 20:0, 20:1, 20:2, 20:3, 20:4, 20:5, 22:0, 22:1, 22:2, 22:4, 22:5 and 22:6 except for TGs which considered: 16:0, 16:1, 18:0, 18:1, 18:2, 18:3, 20:3, 20:4, 20:5, 22:2, 22:3, 22:4, 22:5, and 22:6.

Peak integration was performed with the MultiQuant^TM^ software version 3.0.3. Lipid species signals were corrected for isotopic contributions (calculated with Python Molmass 2019.1.1) and were quantified based on internal standard signals, in adherence to the guidelines of the Lipidomics Standards Initiative (LSI) (level 2 type quantification as defined by the LSI).

### Cathepsin D activity assay

Cathepsin D activity in zebrafish extracts was assayed using a protocol adapted from (Kuster et al. 2019) and (Matsui et al. 2013). Total protein was extracted from a pool of 60 WT and mutant (MUT1) larvae at 5 dpf. Samples were homogenized at 4 °C in 300 μl lysis buffer (0.1 M Na acetate, 1 mM EDTA, 0.1% Triton X-100, pH 4.0) for 60 seconds at 6000 rpm using a Precellys instrument (Bertin Instruments). The homogenates were centrifuged at 4 °C for 30 min at 9300 × *g* and total protein concentration in the supernatants was determined using the Pierce™ BCA Protein Assay Kit (Thermo Scientific).

Cathepsin D activity was assayed in a reaction mixture (175 μl total volume) containing 28.6 mM sodium acetate buffer (pH 4.0), 1.4 % (w/v) hemoglobin, and 20 μg total protein extract. Mixtures were incubated at 37 °C for 3 hours and reactions were stopped by addition of 150 μl of 15 % (w/v) trichloroacetic acid (TCA) followed by centrifugation at 16000 × *g* for 6 min at room temperature. Supernatants (200 µl) were neutralized by addition of 16 µl of 4 M NaOH. The TCA-soluble peptides were measured using the Pierce™ BCA Protein Assay Kit.

### Cerebral organoid derivation

Cerebral organoids were the same as the ones used in Gomez-Giro et al. (Gomez-Giro et al. 2019). They were derived from isogenic control and CLN3^Q352X^ iPSCs, following the protocol described by Lancaster et al. (Lancaster and Knoblich 2014), and extensively characterized in that study(Gomez-Giro et al. 2019),. Cerebral organoids were maintained for 55 days after the beginning of the differentiation (the total duration in culture was 66 days). At the time of collection, organoids were washed once with PBS, snap frozen and stored at -80°C until they were processed. For each genotype, two organoids of four independent organoid derivations were pooled for the metabolomics analysis.

### Electron microscopy methods

Zebrafish larvae at 6 dpf were fixed in 4% glutaraldehyde (ems), 1% paraformaldehyde (ems), and 2.5 mM CaCl_2_ in 0.1 M cacodylate buffer (Sigma) for 1.5 h and post-fixed in 0.1% OxO_4_ for 1h. After dehydration in increasing concentrations of ethanol, the larvae were embedded in agar 100 resin (agar scientific) and cut into 60 nm sagittal sections in two planes, paramedial and at mid-eye level on an ultramicrotome (RMC Boeckeler). Sections were mounted to 2 x1mm copper slot grids (gilder grids) and imaged on a Zeiss Sigma 300 SEM with a STEM detector at an acceleration voltage of 30 kV. For reference of the zebrafish anatomy, the zfin zebrafish atlas by Diever et al. was used (https://zfin.org/zf_info/anatomy.html).

## Data availability

All relevant data has been included in the supporting information. This study includes no data deposited in external repositories.

## Acknowledgements

We are grateful to the fish caretaker team at the LCSB Aquatic Platform for their valuable daily work. We would also like to thank the LCSB Metabolomics Platform for providing technical and analytical support. We acknowledge the use of Biorender.com for Figures 4 (panels A and B) and 8. This work was supported in part by donations from the ATOZ foundation and Mr. Norbert Becker to CLL and by a Pélican award from the Fondation du Pélican de Mie et Pierre Hippert-Faber to UHM. CLL, RRS and ELS acknowledge funding support from the Luxembourg National Research Fund (FNR) for projects C20/BM/14701042 (CLL) and A18/BM/12341006 (RRS, ELS), respectively.

## Author contributions

CLL and ADC conceived and designed the project idea. UHM performed MO experiments, generation of CRISPR mutants and behavioral assays. RSS, CMR, and MW performed metabolite/lipid extractions as well as untargeted metabolomics and lipidomics measurements, data analysis and visualization. CJ and FG developed targeted measurements for GPDs, amino acids and acylcarnitines. AC and Melanie M. contributed to sample preparation for and imaging by TEM. FKB and Michel M. were responsible for scientific input for TEM analysis and interpretation of the TEM results. SP, ADC, JCS, MLCM and ELS gave input for specific parts of the research and overall manuscript preparation. GGG performed the cerebral organoid differentiation. UHM and CLL performed data interpretation and wrote the manuscript in consultation with all other authors.

## Figure S1

**Conservation and expression of the zebrafish *cln3* gene. (A)** Multiple sequence alignment of CLN3 protein sequences shows high conservation across various species. Gaps are indicated with dashes and different shades of blue were used to represent conservation level: dark blue, blue and light blue for 100%, 75% and 50% conservation in the selected sequences, respectively. Point mutations that are known to cause Batten disease in human patients and posttranslational modification sites are highlighted in red and green, respectively. **(B)** Zebrafish *cln3* expression levels at different developmental stages are represented relative to the expression level at 6 hpf. **(C-D)** Zebrafish *cln3* expression levels in the indicated organs are shown relative to the expression levels in the eye. RNA was extracted from the organs of 2-year-old male and female zebrafish for qPCR analysis. All expression levels were normalized to either the *rpl13α* (dark blue) or the *ef1α* (dotted light blue) reference genes. Data shown in panel B are technical replicates of a pool of 30 larvae. Data shown in panels (**C**) and (**D)** are means ± SDs from three biological replicates. hpf, hours post-fertilization.

## Figure S2

**Generation** o**f stable *cln3* mutant lines in zebrafish using CRISPR/Cas9. (A)** Pipeline for the generation of *cln3* homozygous mutants. Cas9 protein was co-injected with *cln3* and s*lc45a2* sgRNAs into one-cell embryos. Chimeric larvae showing a pronounced albino phenotype were raised to adulthood (P0) and outcrossed with *nacre* fish for creating the F1 generation. F1 adult fish were screened for indel mutations in the *cln3* gene. Founders with a predicted early stop codon were in-crossed to create the F2 generation comprising potentially wildtype (+/+), heterozygous (+/-), and homozygous progeny (-/-). **(B)** *cln3* sgRNA was injected into freshly fertilized embryos and genomic DNA was extracted 24 hpf. Agarose gel analysis confirmed the high efficiency of *cln3* sgRNA-Cas9 to induce indel mutations at the target site. **(C)** Left panel: dorsal view of a 4 dpf uninjected control wildtype and a chimeric larva. The albino phenotype can be evaluated in the eye (red arrowhead). Right panel: chimeric male (upper part) and female (lower part) P0 adults. **(D)** Carriers were in-crossed and raised to adulthood. Genotyping of F2 adult fish showed that 46% and 30% of the progeny were homozygous mutants for the non-sense and exon 4 deletion mutation, respectively.

## Figure S3

**In *silico* prediction of transmembrane helices and N-glycosylation sites in huCLN3, zfcln3 and CRISPR mutants.** The TMHMM (v2.0) program predicts 11 putative transmembrane (helical) regions in the wildtype human and zebrafish CLN3 proteins (Moller-Pedersen et al. 2001). In contrast, only one transmembrane domain is predicted for the truncated protein expressed by the MUT1 line, while the protein encoded by the *cln3*^Δex4^ allele (MUT2 line) is predicted to have a shorter luminal loop between the first two transmembrane domains. The NetNGlyc (v1.0) tool predicts four glycosylation sites in human CLN3 (Gupta and Brunak 2002). N49 is inside of a transmembrane domain and therefore unlikely to be glycosylated. In contrast, the zebrafish protein contains five predicted N-glycosylation sites outside of the transmembrane domains. The expression product of the *cln3* null allele (MUT1 line) is predicted to conserve the first four glycosylation sites, whereas the one of the *cln3^Δex4^* allele conserves only the first of these glycosylation sites that is placed in exon 3.

## Figure S4

**PTZ and PTX treatment in cln3 deficient zebrafish larvae. (A)** Heterozygous and homozygous mutant *cln3* larvae (5 dpf) were placed in individual wells of a flat-bottom 96-well plate. Subacute (5, 7.5, and 10 mM) and acute (15 and 20 mM) concentrations of PTZ were added and 10 min after addition of the drug (habituation), the locomotor activity was tracked and the mean velocity was calculated for each condition (40 min). **(B)** The average distance moved within each 2-min time bin under 0 and 10 mM PTZ yields a similar behavioral profile for both genotypes, but higher activity was observed in MUT1 larvae. Error bars represent SEM, plotted only below the means for the sake of clarity (n = 12). **(C)** A similar procedure was performed with PTX as described for PTZ. **(D)** Behavioral profile under PTX exposure (0.3 and 1 mM). Error bars represent SDs, plotted only below the means for the sake of clarity (n = 12). Statistically significant differences between lines were determined using the two-way ANOVA test followed by Dunnett’s multiple comparisons test (****, p ≤0.0001).

## Figure S5

**Untargeted metabolomics analysis using LC-MS in the discovery sample set. (A)** Principal Component Analysis (PCA) analysis for WT (black) and MUT1 (blue) zebrafish larvae based on all variables (*m/z* features) detected in positive and negative modes. The first principal component (PC1) explains 25.7% of the variation, the second component 14.9%, and the third component 10.3%. **(B)** Violin plots showing normalized abundances for the indicated acylcarnitines in WT and MUT1 samples. Statistically significant differences between lines were determined using an unpaired multiple Welch’s t-test (*, p ≤ 0.05; **, p ≤ 0.01; ***, p ≤ 0.001; ****, p < 0.0001).

## Figure S6

**Multivariate and metabolite enrichment analyses of the untargeted metabolomics validation dataset. (A)** Diagram of feature filtering by statistical analysis and compound identification steps. Out of 6373 detected features, 1125 differed significantly between the WT and MUT1 lines and could be annotated. **(B)** PCA analysis using all *m/z* features from the validation set of polar extracts detected in positive and negative modes. The first PCA component (PC 1) already shows segregation of the metabolites from WT and MUT1 samples. Each dot represents a single biological replicate of WT larvae (black) and MUT1 larvae (blue). **(C)** Metabolite enrichment analysis of differential metabolites (p<0.05) highlights the major dysregulated pathways between WT and MUT1 samples in the validation set.

## Figure S7

**Most significantly altered metabolite classes between wildtype and MUT1 larvae in the validation sample set.** Violin plots show normalized abundances for the indicated metabolites values. Statistically significant differences between the two zebrafish lines were determined using an unpaired multiple Welch’s t-test (*, p ≤ 0.05; **, p ≤ 0.01; ***, p ≤ 0.001; ****, p < 0.0001).

## Figure S8

**Indications for impaired proteolytic activity in MUT1 larvae. (A)** Heatmap diagrams from the discovery and validation untargeted metabolomic datasets generated from significantly different oligopeptides containing one prolyl residues. **(B)** CtsD activity assay in WT and MUT1 extracts showing a mild, not statistically significant, decrease of activity in the MUT1 line. **(C)** Western blot analysis of CtsD protein levels in the zebrafish extracts used for the activity assays in panel B. **(D)** Representative electron micrographs of intestine and liver cells of 60 nm sagittal sections of WT and MUT1 larvae at 6 dpf. No fingerprint-like structures were observed in the MUT1 samples. Scale bars represent 3 μm.

## Figure S9

**Analysis of GPDs, and amino acids in polar extracts of WT, MUT1 and atp13a2*^sa18624-/-^* zebrafish larvae using targeted LC-MS analysis. (A)** Representative LC-MS extracted ion chromatograms of GPI (m/z 333,0592; RT 10.92 min) obtained with polar metabolite extracts from WT and MUT1 zebrafish larvae (5 dpf) showing the highest peak at RT 10.92 min. **(B)** GPD analysis in polar metabolite extracts of WT and *atp13a2^Sa18624-/-^* larvae (6 dpf). Each dot represents a pool of ten larvae and in total four biological replicates were measured. **(C)** Amino acids that were not significantly altered in whole larvae extracts of WT and MUT1, based on targeted analysis. Each dot represents a pool of 40 larvae and in total four biological replicates were measured. AA, automatic (integration) area; AH, automatic height.

## Figure S10

**Acylcarnitine measurements using two independent targeted LC-MS methods**. In the upper part, ACs were extracted and measured by Lipometrix (Method 1). In the lower part, carnitine, acetylcarnitine and ACs were extracted and measured at the LCSB metabolomics facility (Method 2). Each dot represents a pool of 40 larvae and in total six (Method 1) and four (Method 2) biological replicates were measured. In both targeted methods, AC showed slightly increased levels in MUT1 compared to WT larvae. Statistically significant differences between the zebrafish lines were determined using an unpaired parametric multiple Wech’s t-test (*, p ≤ 0.05; **, p ≤ 0.01; ***, p ≤ 0.001; ****, p < 0.0001).

## Figure S11

**Targeted analysis of phospholipids in MUT1 larvae.** Stacked bar graphs of fatty acid composition for diverse phospholipid species assayed in nonpolar extracts of 5 dpf zebrafish larvae using targeted LC-MS-based methods. Nonsignificant differences were observed between WT and MUT1 samples for these phospholipids. Each stacked bar is the average of six biological replicates of pools of 40 larvae, normalized against the DNA concentration.

---

**S1 Table: Quantitative and qualitative LC-MS data for metabolomic features in the discovery sample set.**

**S2 Table: Quantitative and qualitative LC-MS data for metabolomic features in the validation sample set.**

**S3 Table: Significant differential metabolic features, revealed by untargeted metabolomics, validated against standard compounds.**

**S4 Table: Quantitative and qualitative LC-MS data for lipidomic features in discovery and validation sample sets.**

**S5 Table: Abbreviations used in this study for the nomenclature of lipid subclasses.**

**S6 Table: Sequences of primers used in this study.**

**S7 Table: Exhaustive list of annotations for metabolomic features detected in the discovery and validation sample sets.**

**S8 Table: Mass transitions and compound-dependent source parameters for targeted method for acylcarnitines and amino acids.**

